# Coarse-graining genome-scale metabolic models for dynamic, multi-scale simulations of *Pseudomonas aeruginosa* in gel environments

**DOI:** 10.64898/2026.11.05.764552

**Authors:** Morgan S. Kim, Jason A. Papin, Roseanne M. Ford

## Abstract

Microbial communities are shaped by coupled interactions between microenvironment conditions and the intracellular adaptations that emerge in response external metabolic cues. For pathogenic microbes such as *Pseudomonas aeruginosa*, these interactions can rapidly generate heterogeneous sub-populations that exhibit adaptations for sustaining host infection and resisting antibiotic treatment. Technological strides in -omics data collection, combined with robust computational tools like genome-scale metabolic models (GEM) for predicting cellular metabolism, have yielded promising multi-scale depictions of microbial population development in simulated spatial environments. These multi-scale model simulations help establish and evaluate mechanistic contexts for how infection-enhancing behaviors might emerge in the population or be influenced by variable perturbations in the biological system. However, the number and rigorous tuning of parameters influencing the connections between intracellular-extracellular events pose barriers when shaping simulations to different *in vitro* systems of interest. In this work, we present a genome-scale, metabolism-informed continuum model (GEM-CM), a computational framework for coupling a reaction-diffusion model to a surrogate GEM representation that generates metabolic rate predictions informed by flux balance analysis outputs. A key feature of GEM-CM is its application of random forest regression, or ensembles, to coarse-grain the interface connecting the GEM and reaction-diffusion models down to a small set of input metabolites methodically chosen through the random forest’s evaluation of feature importance.

To demonstrate GEM-CM’s utility, we evaluated simulation predictions against *P. aeruginosa* PA14 populations radially propagating in low weight percentage agar gels. In addition to biomass growth, we simulated PA14 populations producing the cytotoxic secondary metabolite pyocyanin (PCN). GEM-CM could capture PA14 growth dynamics in agar gels when ensembles were provided with three carbon source inputs describing the simulated growth media. Bacterial motility parameters and the random forest ensemble allowed GEM-CM simulations to accurately predict spatial biomass distributions for PA14 populations over time and probe their PCN production dynamics. Ensembles also demonstrated robust performance for single gene knockout conditions with GEM-CM simulation accurately capturing the partial or total biomass and PCN inhibition observed in transposon mutant screenings. When evaluating transposon mutant population development, the GEM-CM framework additionally suggested that *metH* may be a promising target to inhibit the spatial propagation of PA14. This work’s proof-of-concept applications illustrated GEM-CM utility as an early stage multi-scale modeling tool for interrogating *in vitro* metabolic behavior in microbial populations.

**AUTHOR SUMMARY:** Pathogenic microbial communities alter their environmental conditions as they grow and propagate, which in turn drives the development of heterogenous metabolic behaviors across different spatial regions in the population. Computational simulations using genome-scale metabolic models (GEMs) can offer insights into how bacteria metabolic behaviors can influence larger population-level outcomes. Many of these simulation platforms however involve extensive parameter optimization to situate the GEM to its simulated environment making their application a challenge at the early investigation stages. To address this, we developed a computational approach that innovatively utilizes a random forest regression model to perform metabolic predictions in place of the GEM. This random forest surrogate approach reduces the number of parameters required to connect it to a simulated environment while still providing valuable, informed predictions for bacterial metabolism. We tested this computational approach to simulate *Pseudomonas aeruginosa* propagation and their production of cytotoxic pyocyanin in low weight percentage agar gels. Simulations demonstrated their robustness for capturing biomass and pyocyanin perturbations in *P. aeruginosa* gene knockouts and revealed potential intracellular mechanisms that may inhibit the pathogenic community’s spatial expansion. We believe this multi-scale computational approach provides population models valuable connections to metabolic behaviors, which can inform early stage investigations of crucial variables driving *in vitro* bacterial population development.

## 2. INTRODUCTION

The development and sustained presence of microbial communities are influenced by how cells within the population respond to dynamic fluctuations in their environment niche. As the population propagates through its surroundings via cell growth and other active mechanisms such as flagellar motility, spatial metabolite gradients are generated by microbes consuming local nutrients and producing secondary metabolites that impose heterogeneous environmental stresses across the microbial community [1, 2, 3]. For pathogenic bacterium like *Pseudomonas aeruginosa*, this metabolite heterogeneity can drive extensive intracellular changes that produce sub-populations with adapted metabolic processes specialized for persisting in their local microenvironment and challenging antibiotic treatment efficacy [4, 5, 6, 7]. This scenario is especially hazardous for individuals with cystic fibrosis and chronic obstructive disease as opportunistic *P. aeruginosa* can colonize the obstructive mucus plugs in the lungs and establish chronic, antibiotic resistant respiratory infections that worsen condition prognosis [8, 9, 10]. Thus, initiatives for designing higher efficacy treatments against these bacterial communities are motivated to gather mechanistic understanding of their emergent metabolic heterogeneity in biologically relevant environments [11, 12].

Advancements in -omics technologies, especially high-throughput *in situ* techniques capable of probing molecular markers with single-cell spatial granularity [13], have enabled transformative insights into the diversity of metabolic states within a bacterial community as well as their relation to other metabolic states. Examples of the latter are recent spatial transcriptomic analyses isolating denitrification pathway activation within *P. aeruginosa* aggregates under high oxidative stress conditions [14] and metabolomic studies identifying fatty acid and amino acid exchanges between aggregate interior and peripheral populations during low nutrient availability [15]. One promising computational approach for incorporating this influx of biological data is genome scale metabolic models (GEM), a stoichiometric network representation of metabolic processes informed by an organism’s annotated genome and biochemical databases [16, 17]. Using this network and biological contexts from -omics data, GEMs impose constraints on the ‘flux’ of mass moving through process reactions and can perform flux balance analysis (FBA) that optimizes mass fluxes towards a defined biological objective (e.g. biomass growth). Changes to GEM flux outputs following constraint or network-level alterations can contextualize an organism’s metabolic capabilities under different genomic states or environmental conditions [18, 19, 20, 21]. However, these GEMs in isolation only illustrate an instantaneous perspective of metabolism that is disconnected in the spatial and temporal dimensions.

To address this, several computational frameworks such as BacArena [22], MiMICS [23], MetaBiome [24], ACBM [25], and COMETS [26] have emerged for simulating GEMs as dynamic microbial communities growing within an extracellular environment that tracks the spatial distribution of cells and metabolite resources. COMETS, in particular, can have its extracellular environment scalable to centimeter length scales due to its microbial communities being depicted as bulk, or continuum, populations instead of structures comprised of individual cells [27]. These multi-scale models frequently establish an ‘interface’ between GEMs and the environment where discrete metabolite concentrations are interpreted as GEM flux constraints and FBA outcomes are integrated into partial differential equations (PDE) evaluating metabolite consumption, cell growth, and byproduct generation. Over successive time iterations, the information exchanged at this interface couples intracellular and extracellular events, which enables simulated interactions between metabolic state and local metabolite environment that are valuable for deconvoluting mechanism hypotheses driving -omic trends without extensive lab resource investments. Modular updates to model frameworks, such as COMETS incorporation of multiple cell motility patterns and evolutionary dynamics [27, 28, 29], have allowed users to flexibly tailor these tools to their system of interest; however, tuning model parameters at this GEM-environment interface to recreate relevant microbial community behavior remains a challenge. Variables such as GEM size and spatial discretization can also drastically increase computational time of batch simulations as a result of repeated dynamic FBA calculations for each grid space [29], thus reliance on these tools may be impractical if one is first approaching a population metabolic behavior with minimal *a priori* information.

In this work, we present a multi-scale framework for simulating motile, spatially heterogenous microbial communities using a surrogate model to coarse-grain the computational interface between GEM mechanistic metabolism and the reaction-diffusion model. This approach, referred to as genome-scale, metabolism-informed continuum models (GEM-CM), seeks to train a random forest regression that operates in place of FBA to provide the reaction-diffusion model access to salient metabolic behavior. We hypothesize that random forests can effectively coarse-grain desired GEM predictions (e.g. biomass growth rate and secondary metabolite production) when the regression model training is supported by simulation datasets representing the FBA solution space. Furthermore, the evaluation of training feature importance following random forest regression can offer a methodical approach to reduce the number of input variables informing predictions of bacterial growth and metabolite production in the reaction-diffusion model.

To demonstrate GEM-CM’s utility for probing *in vitro* bacterial populations, we framed model simulations around *P. aeruginosa* PA14 populations in low weight agar gel environments. Surrogate random forest regression models were trained on simulated growth media in order to identify which M9-glucose carbon source metabolites were important to include in the reaction-diffusion model to depict PA14 growth. Alongside PA14 population growth and spatial expansion in agar gel environments, we investigated the production of pyocyanin (PCN), a *P. aeruginosa* secondary metabolite relevant to respiratory infection condition. When secreted by *P. aeruginosa*, PCN has been observed elevating reactive oxygen species generation and oxidative stress in the host lung microenvironment leading to epithelium damage, neutrophil cell death, and respiratory infection exacerbation [30, 31, 32, 33, 34]. By framing different *in silico* PCN synthesis conditions in GEM-CM simulations, we were able to probe prospective PCN production dynamics and contextualized their predictions against observed *in vitro* PA14 dynamics. Lastly, we evaluated GEM-CM capabilities to simulate PA14 under different gene knockout contexts and predict potential growth or PCN synthesis inhibition relative to wild type PA14. GEM-CM predictions were then validated against swim plate assay screenings of single-insertion transposon mutants. Overall, we believe these applications illustrate GEM-CM’s promising role as an early computational tool, preceding techniques like COMETS, for probing metabolic behavior in spatially heterogeneous microbial communities.

## 3. RESULTS & DISCUSSION

### 3.1: Genome-scale, metabolism-informed continuum models (GEM – CM) for simulating spatially heterogenous bacterial communities and their metabolic activity

The multi-scale computational framework GEM-CM was developed to simulate the spatial expansion and metabolic capabilities of PA14 communities while also simplifying the information exchanged between the GEM and its continuum environment down to a small set of salient metabolites (**Fig 1**). The continuum layer tracking PA14 movement, metabolite diffusion, and metabolite consumption drew its foundation from a 1-D reaction-diffusion model established in Kim *et al.* for describing PA14 communities radially expanding in a stationary environment from an inoculation point [35]. Secondary metabolite generation was added to the model in order to simulate the community’s production of cytotoxic PCN. Reaction rates for PA14 growth, nutrient consumption, and PCN production in the continuum layer are informed by a random forest (RF) regression model, referred here as an *ensemble*, which operates as a surrogate to FBA calculations for metabolic behaviors of interest. Similar machine learning surrogate models showcased by Song *et al.* demonstrated robust predictions of carbon source utilization shifts in *Swewanella oneidensis* using limited metabolite inputs and showed reduced computational costs compared to simulations of equivalent spatial and temporal discretization that directly performed FBA [36]. Additionally, RFs have been applied as classification models for GEM analysis to help isolate metabolic pathways and reactions that were crucial to bacteria expressing highly pathogenic phenotypes [37, 38]. Here, we sought to leverage RFs to both methodically identify carbon source variables for the continuum layer via feature importance scoring and train ensemble models to predict desired FBA outputs that inform PDE reaction terms.

**Fig 1.**
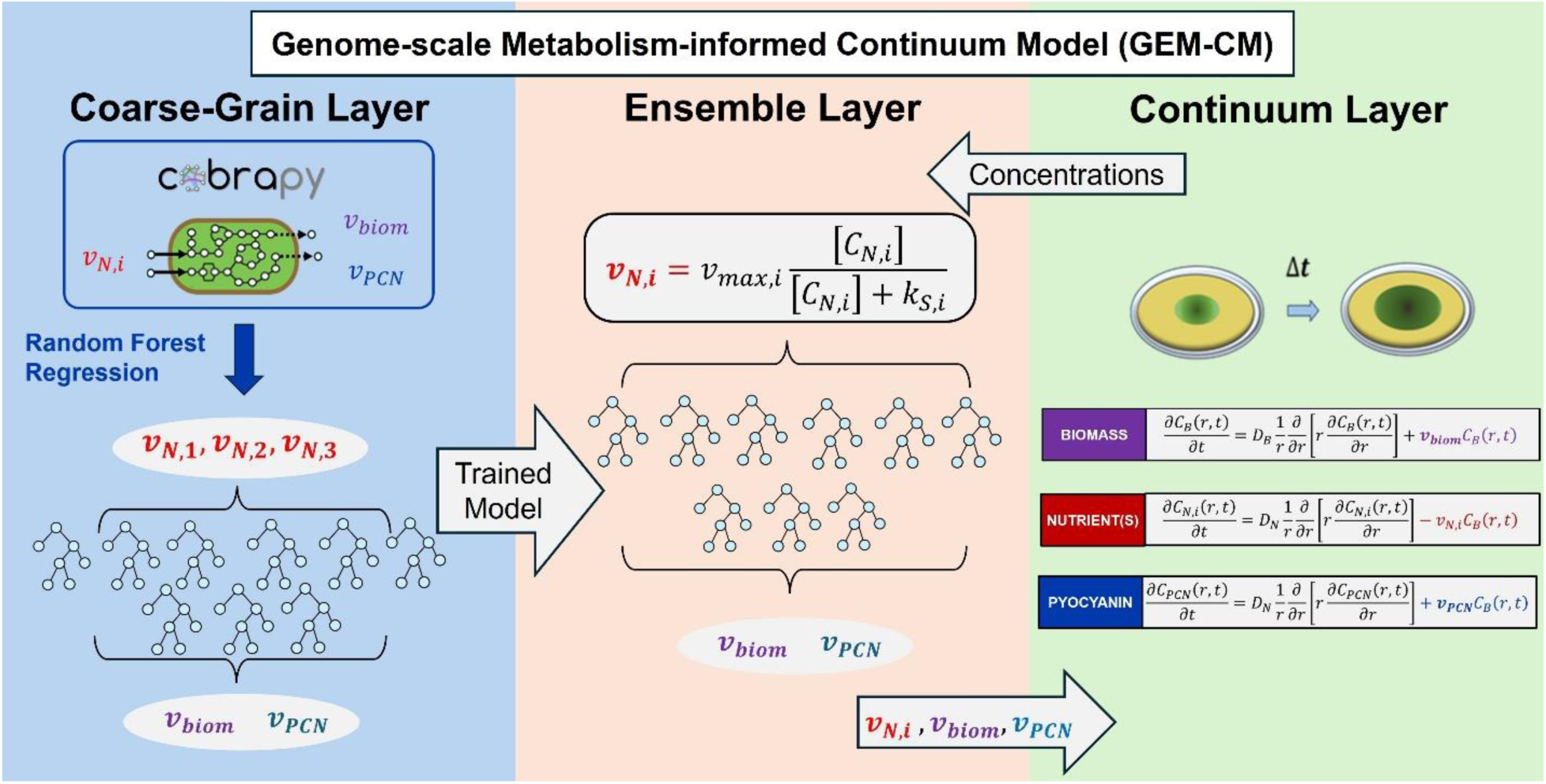
Flow chart representation of the genome-scale metabolism-informed, continuum models (GEM-CM). PDEs in the continuum layer describe the spatial flux and bacteria generation over time in the swim plate along with nutrient consumption and PCN production. For each discretized space in the continuum layer, the trained ensemble receives metabolite concentration(s) as reaction flux inputs and returns growth, consumption, PCN synthesis rates to the continuum layer. The ensemble is prepared by the coarse-grain layer that trains a random forest regression on FBA outputs generated from a GEM of the microorganism.

To first build the RF training foundation for ensemble models, GEM-CM compiled FBA simulation datasets of biomass and PCN synthesis fluxes generated by a GEM of iPau21, a *P. aeruginosa* PA14 genome-scale metabolic network reconstruction [20]. We narrowed the FBA solution space by only varying the flux constraints of GEM exchange reactions associated with major carbon sources in chemically defined M9-glucose growth media (**Table S.3**) in order to emphasize external nutrient substrate contributions to biomass flux. When passing all the varied exchange reaction flux constraints as input features and setting biomass flux as the prediction target, GEM-CM routines in the coarse-grain layer revealed that a majority of feature importance was concentrated in the exchange reactions for L-arginine, L-glutamate, and glucose. The final model used in the ensemble layer was an RF regression trained using only the L-arginine, L-glutamate, and glucose exchange reactions in the FBA simulation dataset to predict biomass and PCN synthesis fluxes. Ensemble hyperparameters and prediction accuracy against hold-out FBA data are presented in **Table S.7** and **Fig S.2**, respectively.

The metabolites transported by the final ensemble’s training exchange reactions were defined as GEM-CM carbon sources. During GEM-CM simulations, discrete concentrations from the continuum layer are passed to the ensemble layer which then updated reaction rates (*v_biom_*, *v_prod_*, *v_cons,glucose_*, *v_cons,L-Arg_*, and *v_cons,L-Glu_*) according to their ensemble-predicted values. These updates occurred at five minute intervals to mimic the approximate time scale for mRNA translation in the cell to respond to extracellular fluctuations [39]. To translate metabolite concentration into ensemble layer mass flux inputs, we applied a Monod equation relationship (**Eq. 1**) previously used by other dynamic FBA frameworks [40, 41, 42].

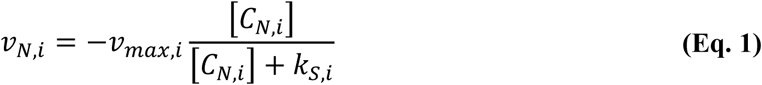

Through **Eq. 1**, the ensemble inputs (*v_N,i_*) are bound by the maximum exchange flux (*v_max,i_*) and varies with external concentration (*C_N,i_*) when metabolite abundance approaches the half-velocity constant (*k_S,i_*). Each GEM-CM carbon source’s *v_max,i_* and *k_S,i_* were presumed to be unique, and appropriate values were optimized using simulation tests that scored predicted metabolite consumption against *P. aeruginosa* liquid culture data reported by McGill *et al.* [43]. Iterative fitting of *v_max,i_* and *k_S,i_* applied Bayesian optimization to minimize residual error between predicted and measured concentrations for glucose, L-arginine, and L-glutamate (**Fig 2**). The final *v_max,i_* and *k_S,i_* values used for each carbon source were derived from the geometric means of the 10 best-fitting candidates after multiple Bayesian optimization instances (**Table S.10**). Following parameter optimization, GEM-CM simulations were prepared for evaluation against *in vitro* swim plate motility assays.

**Fig 2.**
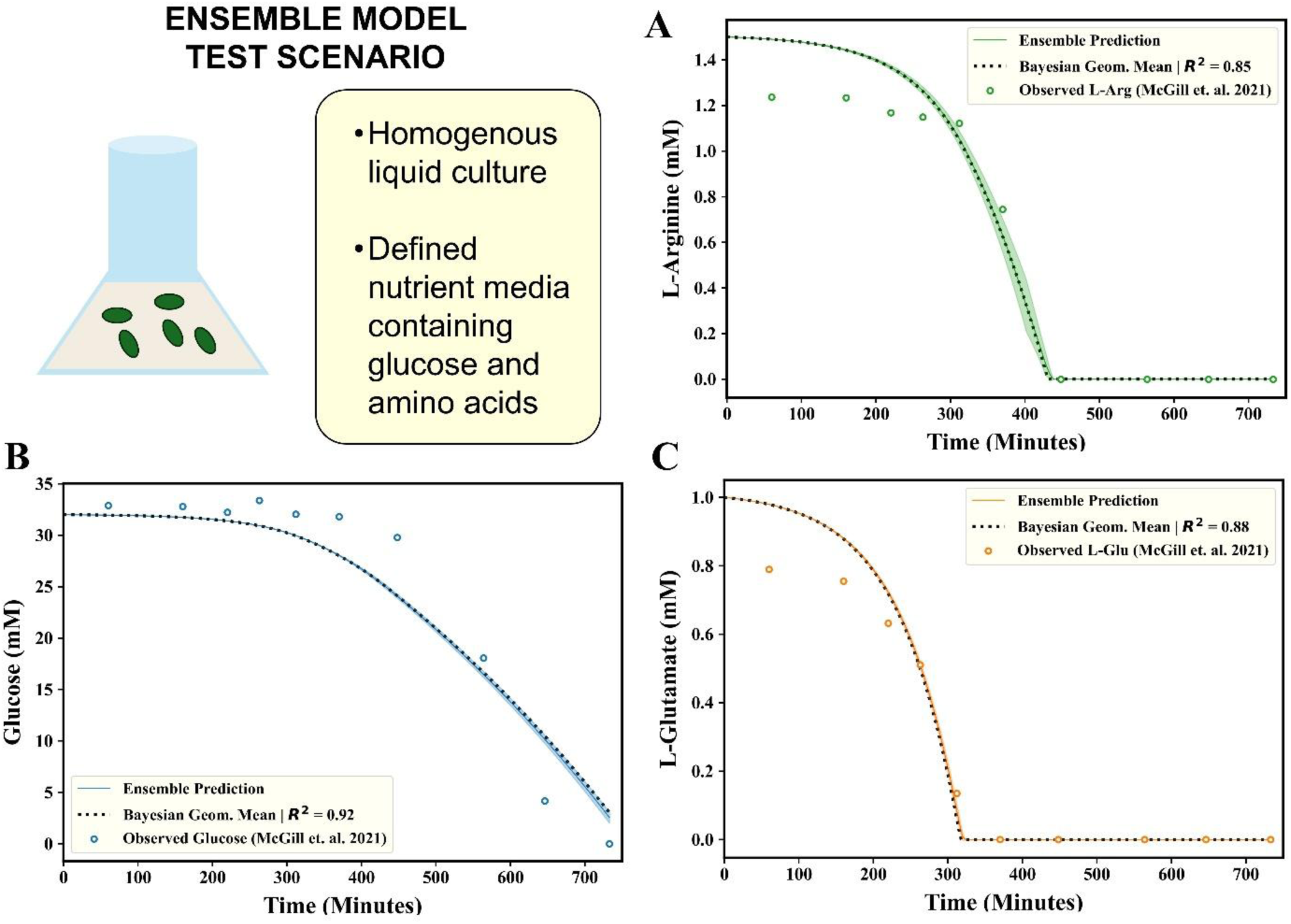
Ensemble model performance versus experimental measurements. Ensemble dynamic simulations model an agitated liquid culture with (**A**) L-arginine, (**B**) L-glutamate, and (**C**) glucose concentrations plotted over time. Average ensemble predictions from the top Bayesian optimization candidates (n = 10) are presented as solid lines with shaded areas representing 1 standard deviation. Ensemble predictions using the *v_max_* and *k_S_* geometric means of Bayesian candidates are presented as a dashed line with its coefficient of determination with concentration data. Liquid culture metabolite data was curated and reported by McGill *et al.* 2021 [43].

### 3.2: GEM-CM performance for predicting swim plate assay biomass distribution and PCN production

For swim plate assays, PA14 was inoculated at the center of a well dish filled with a thin 0.25% w/v agar gel supplemented with M9-glucose growth media. Over time, active bacterial motility and PA14 growth supported by M9-glucose media establish a large bacterial community, referred to as a *macrocolony*, that expands outward towards the well dish wall (**Fig 3A**). The macrocolony development in swim plates (n = 5) was monitored using a widefield microscope to collect tiled images of the well dish over one hour intervals. Simulations (n = 5) were initialized with identical starting carbon source concentrations (**Table S.3**), initial biomass density (grams dry weight/L, gDW/L), ensemble model, and **Eq. 1** parameters. However, each GEM-CM batch simulation was assigned a unique bacterial diffusivity coefficient (*D_B_*) for swimming motility. Repeating practices established in Kim *et al.* [35], *D_B_* were empirically estimated from the macrocolony spreading velocities (**Fig 3B**, **Table S.5**) and the Fisher-KPP traveling wave equation (**Eq. 2**) [44, 45] substituting *µ_max_* with the ensemble maximum biomass flux (*v_biom_*) prediction for M9-glucose media. For 0.25% agar swim plates, *D_B_* values ranging from 0.9 – 1.6E-06 cm^2^/s (**Table S.9**).

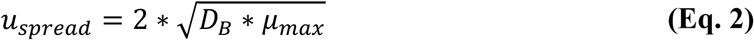

GEM-CM simulations were evaluated over 20 hours, and model results were analyzed for bulk average PA14 growth dynamics and average radial biomass density profiles to compare against experimental observations (**Figs 3C - F**). To compare experiment and simulation data over the same basis, bulk and localized biomass density quantities are scaled to the average endpoint biomass density (*C̅_B,Final_*), which for experiments was equivalent to the average swim plate gDW/L measured at t = 20 hours (**Table S.5**) and equivalent to the density profile bulk average at a specific time point *t_sim_* for GEM-CM predictions (**Eq. 3**).

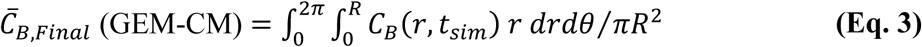

**Fig 3.**
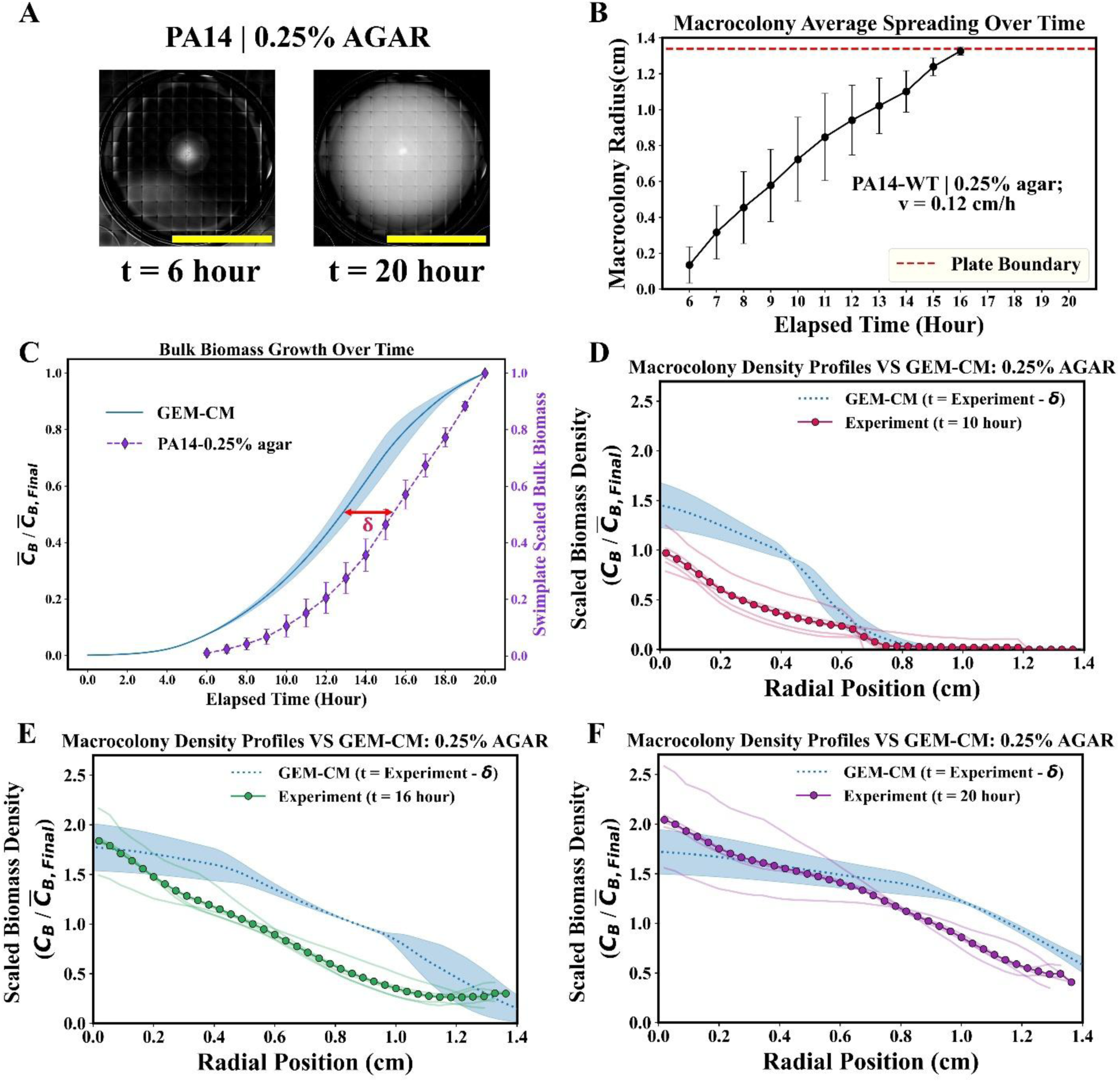
Swim plate assay test scenario for GEM-CM. (**A**) Representative tiled images a PA14 macrocolony (opaque white regions) propagating outward from its central inoculation point. (**B**) Average macrocolony radii versus time and mean PA14 spreading velocity in 0.25% agar swim plate assays (n = 5). Error bars represent 1 standard deviation and the dashed red line denoting the distance to the impermeable well dish wall. (**C**) Average scaled biomass density in experimental and GEM-CM simulated swim plate assays over time. Shaded area around the GEM-CM average represents 1 standard deviation. (**D - F**) Experimental and GEM-CM predicted biomass density profiles at discrete time points scaled to each group’s average endpoint biomass density. GEM-CM density profile predictions are adjusted by the time lag **δ** determined from (**C**). Measured swim plate assays have their group average biomass density visualized as a profile of connected solid markers and sample replicate density profiles visualized as thinner line plots surrounding the average profile.

When first comparing scaled bulk biomass data from swim plate experiments to GEM-CM at matching simulation time points, we observed an apparent time delay (**δ**) of approximately two hours in experimental macrocolony growth dynamics relative to GEM-CM predictions (**Figs 3C**). This **δ** was interpreted to be a growth lag phase experienced by PA14 during swim plate incubation, and all GEM-CM predictions comparisons were accordingly adjusted by –**δ** hours in simulated time (i.e. *C̅_B,Final_* for GEM-CM is evaluated at *t_sim_* = 18 hours). Following simulated time adjustment, spreading distances of the leading macrocolony edge over time exhibited fair agreement between predicted biomass density profiles and experiment profiles suggesting that **Eq. 2** can provide a quick empirical estimation of *D_B_* (**Figs 3D-F**). GEM-CM simulations did exhibit a tendency for overpredicting local biomass density along the macrocolony radial profile, most notably at earlier time points for regions near the macrocolony center. One model component that might have driven these overpredictions is FBA inflating biomass flux / growth rate estimations. While FBA optimization of biomass flux objectives is expected to strongly idealize metabolic capabilities, we also suspect that simulated growth is enhanced by the PA14 GEM being able to simultaneously utilize all the carbon source mass fluxes made available by FBA constraints. Studies extensively monitoring *P. aeruginosa* carbon utilization behavior have reported that the pathogen consumes carbon sources in a sequential order such that glucose is consumed last after other amino acids and organic acids are depleted [46, 47, 48]. Although more tailored FBA objectives could be applied in the future to improve accuracy at early time points, our initial framework for macrocolony growth in M9-glucose did demonstrate improved biomass profile accuracy over longer timescales as seen with endpoint GEM-CM predictions having R^2^ = 0.74 to the t = 20 hour swim plate profiles.

In addition to biomass density distribution predictions, we investigated GEM-CM’s application for probing PCN production behaviors in spatially heterogeneous PA14 communities. To vary PCN synthesis in ensemble models, we framed GEM FBA simulations of PA14 as capable of transitioning between two objectives depending on *in silico* growth condition: (1) maximizing biomass flux or (2) maximizing the sum of biomass and PCN fluxes. When evaluating each set of flux constraints in the FBA simulation training dataset, the objective function during FBA was selected through a decision tree process (**Fig 4A**) based on rules established around the biomass flux. Generally, non-zero PCN synthesis was active if flux constraints produced biomass flux after FBA simulation that could be binned between a feasible minimum (0.05 hour^-1^) for concurrent growth and PCN production and a threshold value (**Γ**). FBA instances with biomass flux that did not meet the feasible minimum or exceeded **Γ** did not synthesize PCN.

**Fig 4.**
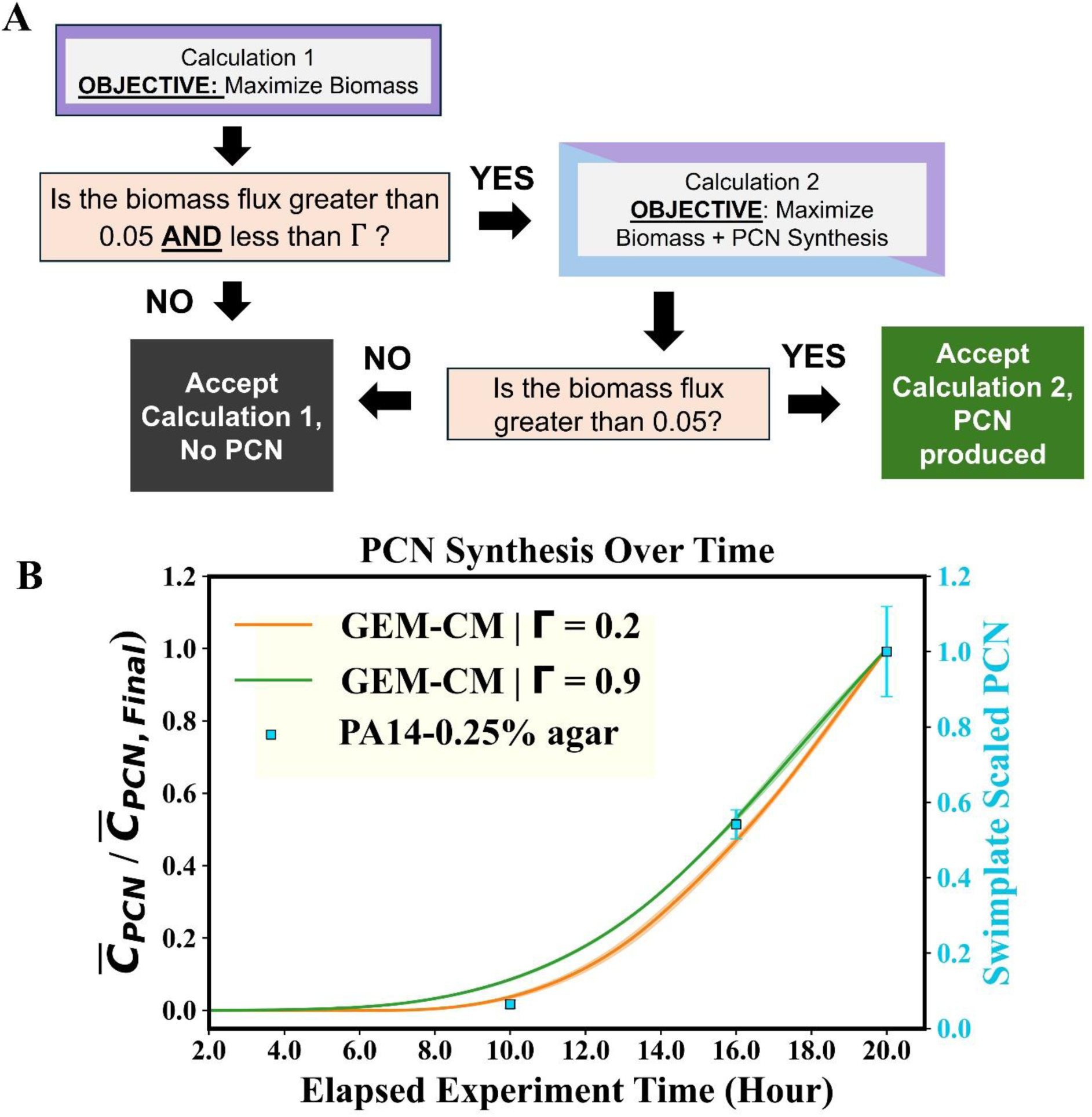
GEM-CM simulations for characterizing PCN production in PA14 macrocolonies. (**A**) Decision tree for selecting FBA objectives based on rules structured around biomass flux. Both FBA simulations ‘Calculation 1’ and ‘Calculation 2’ are performed with identical exchange reaction constraints. (**B**) Scaled PCN concentration in the bulk swim agar or simulation space over time. GEM-CM simulations (n = 5) were performed with **Γ** = 0.2 (orange) and **Γ** = 0.9 (green) ensembles while sharing all other parameter settings; standard deviation across predictions are visualized as shaded areas around the line plot(s). Simulated time is adjusted by ***δ*** such that the simulation endpoint at *t_sim_* =18 hour is matched to t = 20 hour. Three sets of swim plate experiments (batches of n = 3, cyan squares) were measured for PCN concentrations at 10, 16, and 20 hours after PA14 inoculation (**Table S.4**).

As a proof-of-concept test, we trained two ensemble models with FBA datasets that portrayed PCN either being produced regularly by PA14 (Γ = 0.9 hour^-1^) or only produced by PA14 during slower growth conditions (Γ = 0.2 hour^-1^) (**Fig S.3**). GEM-CM simulations (n = 5) were performed for each Γ-ensemble with identical parameter settings and *D_B_* values as the batch simulations used for **Fig 3**. Batch swim plate assays (n = 3) were conducted such that macrocolonies at 10, 16, and 20 hours were measured for bulk PCN concentrations. Scaling PCN concentration from both experiments and GEM-CM by their respective endpoint averages, PCN accumulation in simulated swim plate assays is visualized in **Fig 4B** against relative PCN measured at the sampling time points. Experiments had the majority of PCN production from PA14 occurring between 10 and 20 hours of swim plate incubation. GEM-CM results also aligned with ensemble training expectations with PCN emerging earlier in the Γ = 0.9 simulations than Γ = 0.2 simulations. Interestingly, PCN dynamics within each ensemble group exhibited little perturbations to the tested range of *D_B_* values (0.9 – 1.6E-06 cm^2^/s), which might suggest that bacterial motility variance to this extent does not influence *in silico* growth conditions in the continuum layer. Although normalizing PCN concentrations may occlude distinctions between Γ-ensembles at later time points, we interpreted the better prediction accuracy of Γ = 0.2 simulations for relative PCN at t = 10 hours as an indication that PA14 macrocolonies likely produced PCN under a conditional growth context rather than a continuous production context. This characterization for PCN production by PA14 aligns with previous studies reporting PCN synthesis in *P. aeruginosa* cultures being initiated near the stationary growth phase [49, 50].

### 3.3: GEM-CM simulations capture essentiality of genes closely associated with growth and pyocyanin synthesis

When considering the major role FBA has during the ensemble training process, we were motivated to evaluate GEM-CM performance for gene knockout (KO) scenarios. Gene-protein-reaction rules pairing genes to metabolic pathways in the GEM allow FBA simulations under contexts of gene KO states [16, 51]. Objective perturbations after gene KO, if any, then guide interpretation over the gene’s essentiality in the metabolic activity of interest. For the scope of this investigation, we selected a small pool of gene KO candidates based on high confidence in their biomass growth or PCN synthesis perturbations. Two candidates, ΔargG and ΔpurH, were selected for their reported poor growth fitness in M9 media during PA14 KO screenings conducted by Poulsen *et al.* [52]. Two other candidates, ΔphzM and ΔmetH, were selected from an *in silico* single gene deletion screen performed using iPau21 and an FBA objective fixed to PCN synthesis. Upon referencing gene annotations in the KEGG database, we also suspected ΔmetH had growth fitness reduction arising from disruptions to L-methionine metabolism.

The PA14 KO strains for swim plate assay experiments originated from a transposon insertion library resource previously developed by Liberati *et al.* [53]. M9-glucose swim plate assays were prepared again with an inoculum of WT or transposon mutant PA14 strain and incubated for 20 hours. Swim plates were then imaged for macrocolony biomass density distributions (**Fig 5**) and processed for endpoint bulk biomass density and PCN concentration measurements (**Table S.4**).

**Fig 5.**
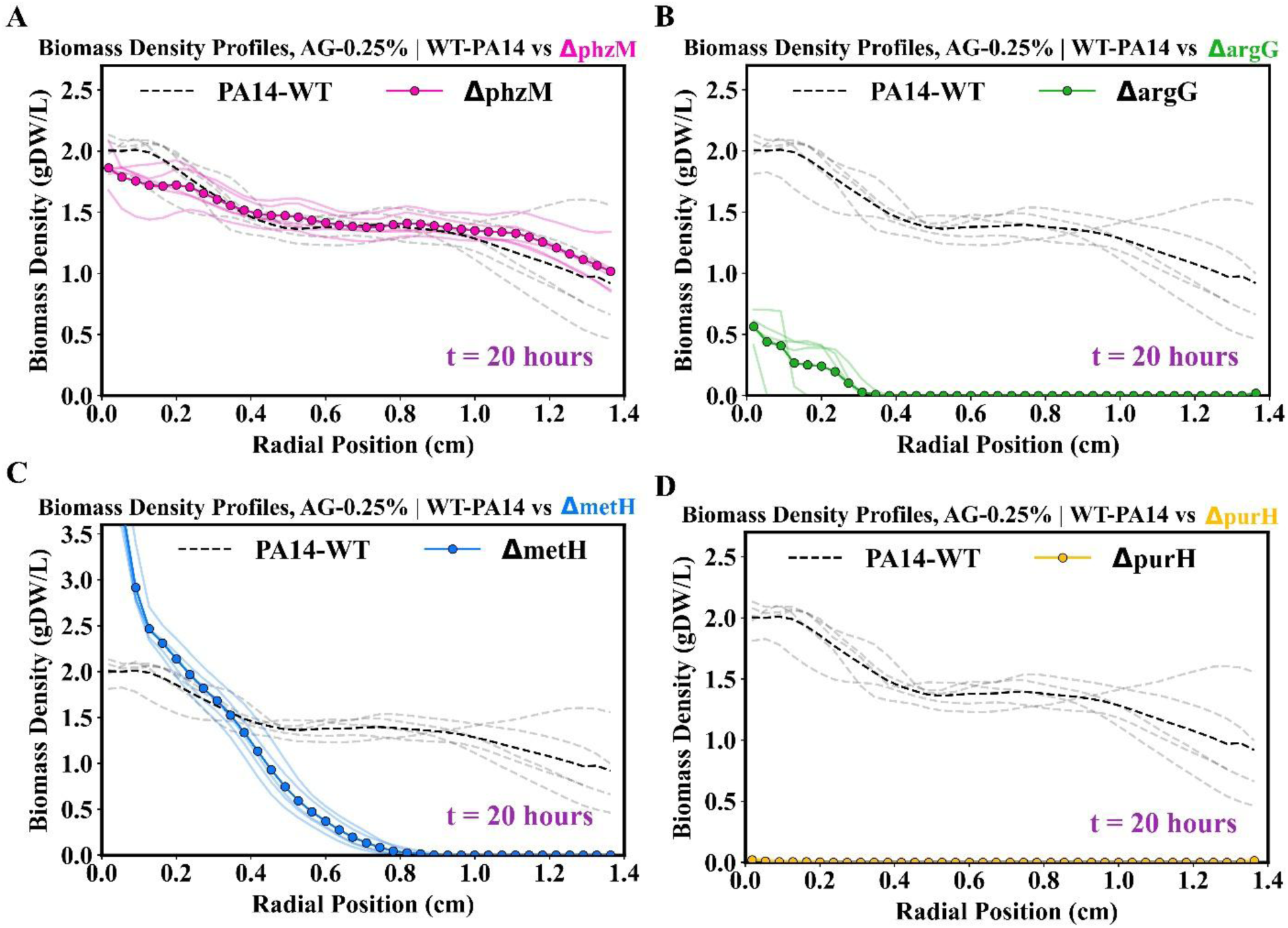
Biomass density profiles of PA14 transposon mutant biomass versus wild type. Biomass density profiles measured after 20 hours of swim plate assay incubation. Group average and sample replicate density profile visualization follow the schemes from Fig 3. ΔphzM **(A)**, ΔargG (**B**), ΔmetH **(C)**, and ΔpurH (**D**) samples (n = 5) are plotted against the same set of wild type swim plate assay data (n = 5).

Overall, transposon mutant biomass density profiles matched the PA14 growth disturbances expected for each KO group. ΔphzM macrocolonies were observed with average biomass density profiles similar to WT macrocolonies (**Fig 5A**) suggesting that *phzM* KO did not inhibit how the strain could grow and spread through 0.25% agar. In contrast, the biomass density profile reduction observed in the ΔpurH, ΔargG, and ΔmetH assays (**Figs 5B-5D**) pointed to each group’s respective gene KO disrupting some metabolic pathway crucial for PA14 growth, which all align with the reported biological roles of *purH*, *argG*, and *metH* genes. The enzyme from *purH* transcription participates in the biosynthesis of adenosine monophosphate and guanosine monophosphate purine nucleotides necessary for sustaining cell growth and viability [54, 55, 56]. Swim plate observations also support the overall growth essentiality of this pathway with ΔpurH samples having significantly (*p* < 0.001) lower biomass density than the second lowest KO group, ΔargG. The genes *argG* and *metH* have their transcribed enzymes participating in arginine and methionine biosynthesis pathways, respectively, and help the cell sustain steady intracellular supplies of amino acids [57, 58].

To evaluate GEM-CM capacities to simulate KO PA14 populations and predict emergent biomass growth or PCN production inhibition from KOs, we repeated high-importance carbon source selection and ensemble model training processes on FBA datasets generated from a deleted gene context. Aside from target gene deletion, FBA simulations retained the Γ = 0.2 hour^-1^ decision tree rule and applied the same exchange reaction constraints used to sample the WT FBA solution space. We note that ΔpurH FBA simulations produced monotonic biomass and PCN synthesis fluxes making the ΔpurH ensemble arbitrarily possess the same training features as the WT ensemble. Carbon source selections did not fluctuate for ΔphzM and ΔargG conditions while ΔmetH had elevated the importance of L-methionine (**Table S.7**). This change in carbon source importance may have been produced by ΔmetH FBA simulations having a greater demand for extracellular L-methionine as its L-methionine biosynthesis pathways were cut off by *metH* inactivation. When restricting input feature training to each KO condition’s three major carbon source exchange reactions, ΔphzM, ΔargG, and ΔmetH ensembles all had a root mean squared error (RMSE) for hold-out biomass flux data near the WT ensemble RMSE (**Fig S.2**). GEM-CM simulations ran with KO ensembles repeated the *v_max_* and *k_S_* parameters for glucose, L-arginine, and L-glutamate as well as *D_B_* values used by simulation runs in the previous section. For ΔmetH simulations, fitted *v_max_* and *k_S_* parameters for L-methionine were applied (**Table S.10**). To emphasize the observed and predicted KO impact to growth and PCN production, GEM-CM predictions were compared to swim plate assay bulk measurements such that simulations and experimental KO endpoint (t = 20 hours) averages were scaled by average WT biomass density or PCN concentrations (**Fig 6**).

**Fig 6.**
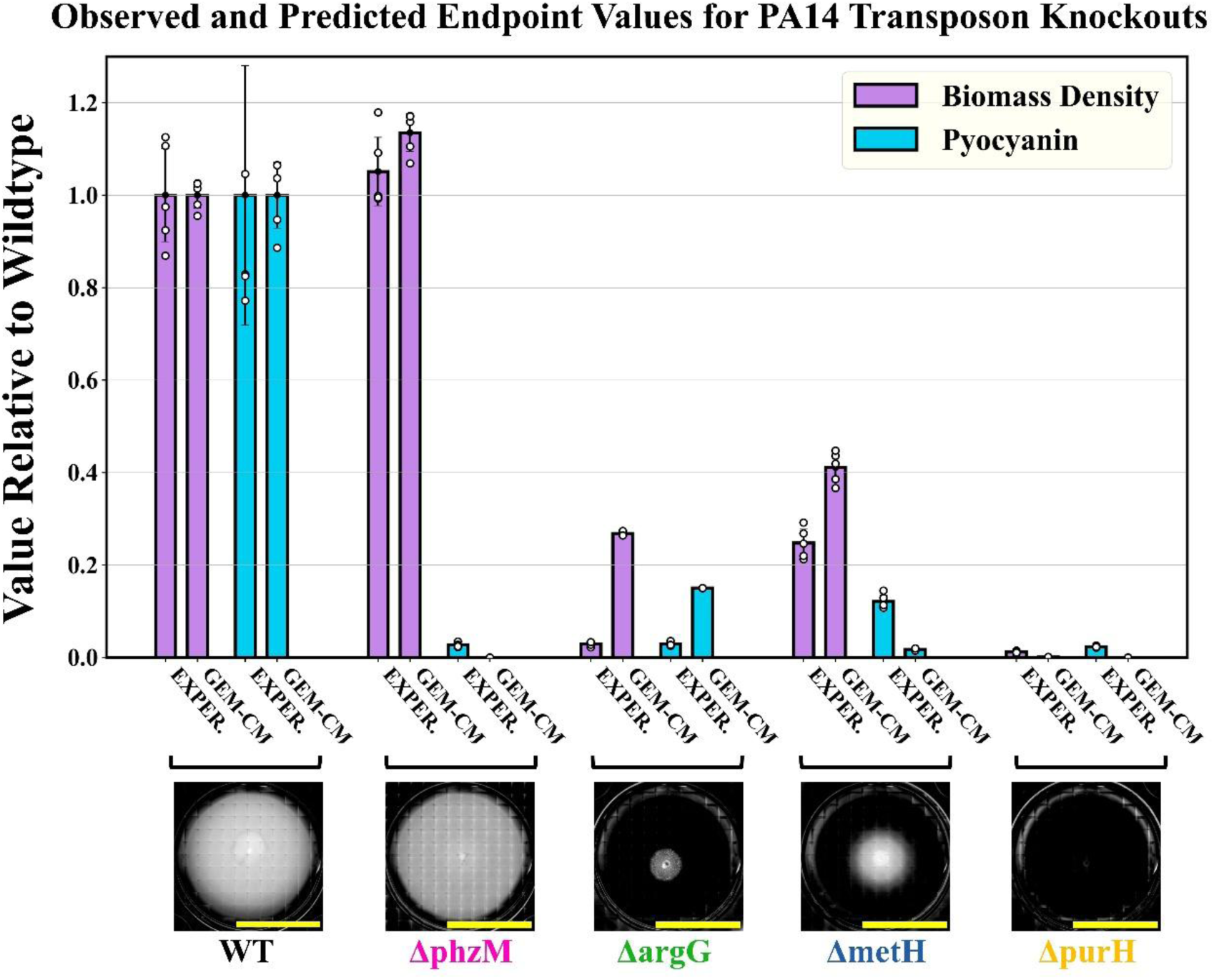
Relative biomass density and PCN from swim plate assays and GEM-CM simulations. For each bar cluster, the first and third columns depict experiment observations and GEM-CM results are depicted by the second and fourth columns. Bar chart heights convey the average endpoint quantities of experiments and simulations relative to the wild type (WT) group with error bars representing 1 standard deviation. Scatter points over each bar depict each sample/simulation replicate value. Cluster labels are accompanied by a representative swim plate assay image, scale bar = 2 cm.

GEM-CM simulation predictions for each KO group were accurate within the context of whether the gene KO would inhibit macrocolony biomass growth or PCN production. The relative KO biomass growth inhibition predicted by ΔargG and ΔmetH simulations were both more moderate compared to their experimental counterparts. Isolated probing of ΔargG/ΔmetH ensembles prediction spaces at low L-arginine/L-methionine levels against a non-mutant ensemble did confirm KO conditions returning notably lower biomass flux compared to non-mutant predictions (**Figs S.5 & S.6**). However, ensemble prediction spaces at high L-arginine or L-methionine exhibited little apparent differences between KO and non-mutant conditions. Overall, we interpreted the cases of biomass flux inhibition in ΔargG and ΔmetH models as supporting the ensemble approach’s capabilities to identifying the altered amino acid biosynthesis mechanisms in the GEM and retain these metabolic changes throughout coarse-graining and regression training. Given that L-arginine / L-methionine was initially available in the environment media for ΔargG / ΔmetH GEM-CM simulations, our estimated amino acid consumption rates combined with inflated ensemble growth rate predictions could have led to an extended time window for KO PA14 growth *in silico* sustained by lingering amino acid concentrations prior to total depletion.

ΔpurH and ΔphzM macrocolony simulations successfully predicted total KO inhibition of PA14 growth PCN production, respectively, with ΔphzM also having predicted no growth inhibition. However, we believe these GEM-CM performances are biased by *purH* and *phzM* having direct influence over these PA14 metabolic capabilities through gene-protein-reaction rules that are situated at the bottleneck points of reaction pathways. In the GEM reaction network, *purH* can alter nucleotide availability in the reaction for biomass flux and *phzM* allows the synthesis of an intermediate molecule that is upstream of PCN [59, 60]. These genes’ direct influence is further illustrated by the ΔpurH ensemble being unable to output non-zero biomass flux predictions and the ΔphzM ensemble being unable to predict non-zero PCN synthesis flux (**Fig S.4**). Interestingly, both ΔphzM experiment and GEM-CM groups had greater average biomass at the 20 hour endpoint than the WT group, though this difference was not statistically significant for the swim plate experiments. From an FBA simulation perspective, cutting off secondary metabolite pathways could allow more mass flux through the biomass generating reactions. PCN has also been reported to have self-cytotoxic mechanisms in *P. aeruginosa* communities during competitive conditions for local environment resources [61, 62]. This could have made the ΔphzM knockout more unbounded in its growth without the cytotoxic inhibition of PCN exposure, though future investigation would need to separate the metabolic burden of PCN synthesis versus PCN exposure to better address the larger biomass density from ΔphzM swim plate assays.

For both swim plate assay and GEM-CM simulation cases, KO groups that experienced biomass growth inhibition also had lower endpoint PCN relative to WT. However, ΔmetH macrocolony simulations were distinguishable from other KO groups due to their pronounced reduction in PCN production. Isolated probing of the ΔmetH ensemble also determined that PCN synthesis fluxes predictions were nearly zero across the range of input carbon source fluxes (**Fig S.6**). In searching literature, we were unable to identify a prior study directly linking *P. aeruginosa metH* knockout and PCN production, yet mechanisms in the KEGG database have led us to hypothesize that *metH* influences PCN through S-adenosyl-methionine (SAM). L-methionine synthesized by the *metH-*associated enzymatic reaction feeds into SAM synthesis, which then participates as a co-factor for methyl group transfer reactions like the *phzM* enzyme reaction [60] (**Fig 7**).

**Fig 7.**
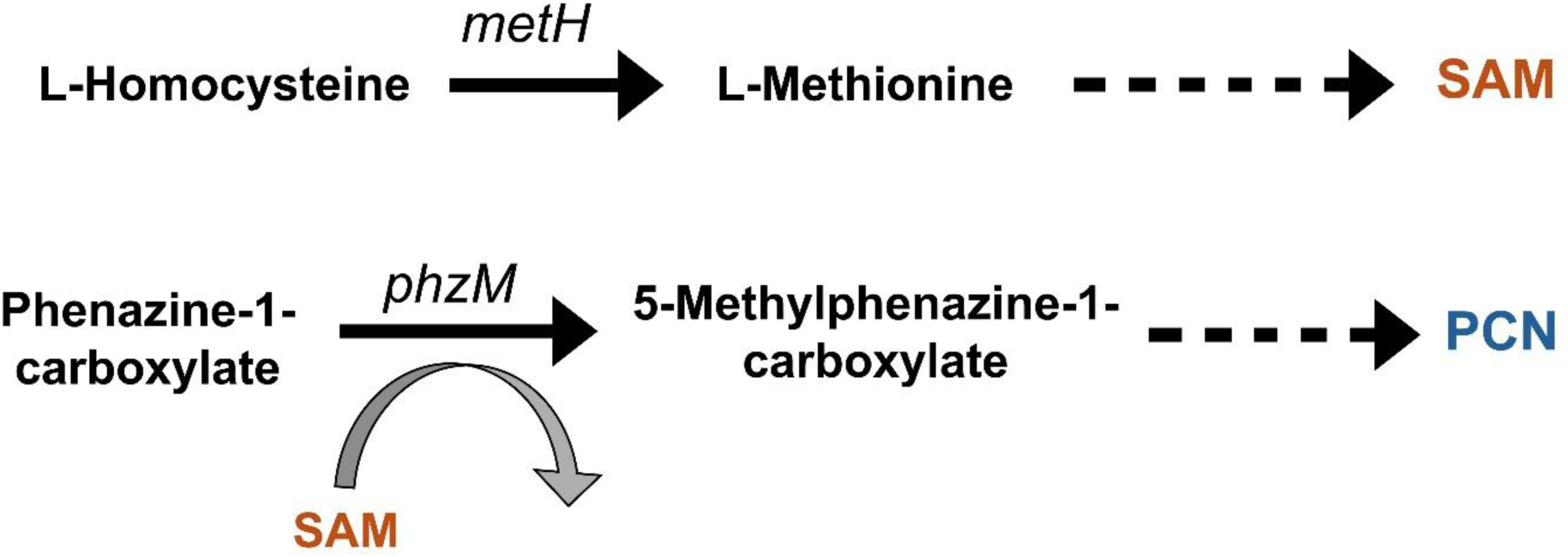
*metH* role in PCN synthesis. Simplified enzyme-reaction pathways for SAM and PCN synthesis emphasizing the pathway points that involve *metH*, *phzM*, or SAM.

Efforts to validate this mechanism for SAM’s involvement in PCN synthesis through flux variability analysis are presented in further detail in *Supplementary Materials, Appendix G*. To briefly highlight analysis results, we observed decreased upper flux bounds for SAM generation and PCN synthesis reactions in iPau21 after *in silico* deletion of *metH* confirming that SAM limitation and PCN flux reduction occurred concurrently in the ΔmetH GEM context (**Table S.11**). Although ΔmetH swim plate samples also had significantly lower (*p* < 0.01) mmol PCN per gDW biomass that the WT group, we were unable to interpret if this PCN yield reduction exclusively driven by this hypothesized mechanism due to the underlying growth deviations also exhibited by ΔmetH macrocolonies.

### 3.4: metH knockout decreases PA14 bacterial motility in swim agar

The unique properties of high local biomass density and short spreading distance observed in ΔmetH density profiles (**Fig 5C**) continued to draw our interest around the KO beyond its growth inhibition. Biomass density near the center of ΔmetH macrocolonies greatly exceeded the biomass densities of ΔargG and ΔpurH samples indicating that ΔmetH had better growth fitness in M9-glucose agar. ΔmetH macrocolonies also spread less than one centimeter from their inoculation point after 20 hours of incubation suggesting that the KO might have had confined macrocolony expansion. To confirm these trends, we collected WT and ΔmetH M9-glucose growth curves and performed timelapse imaging on a second set of ΔmetH swim plates (**Figs 8A-8B**).

**Fig 8.**
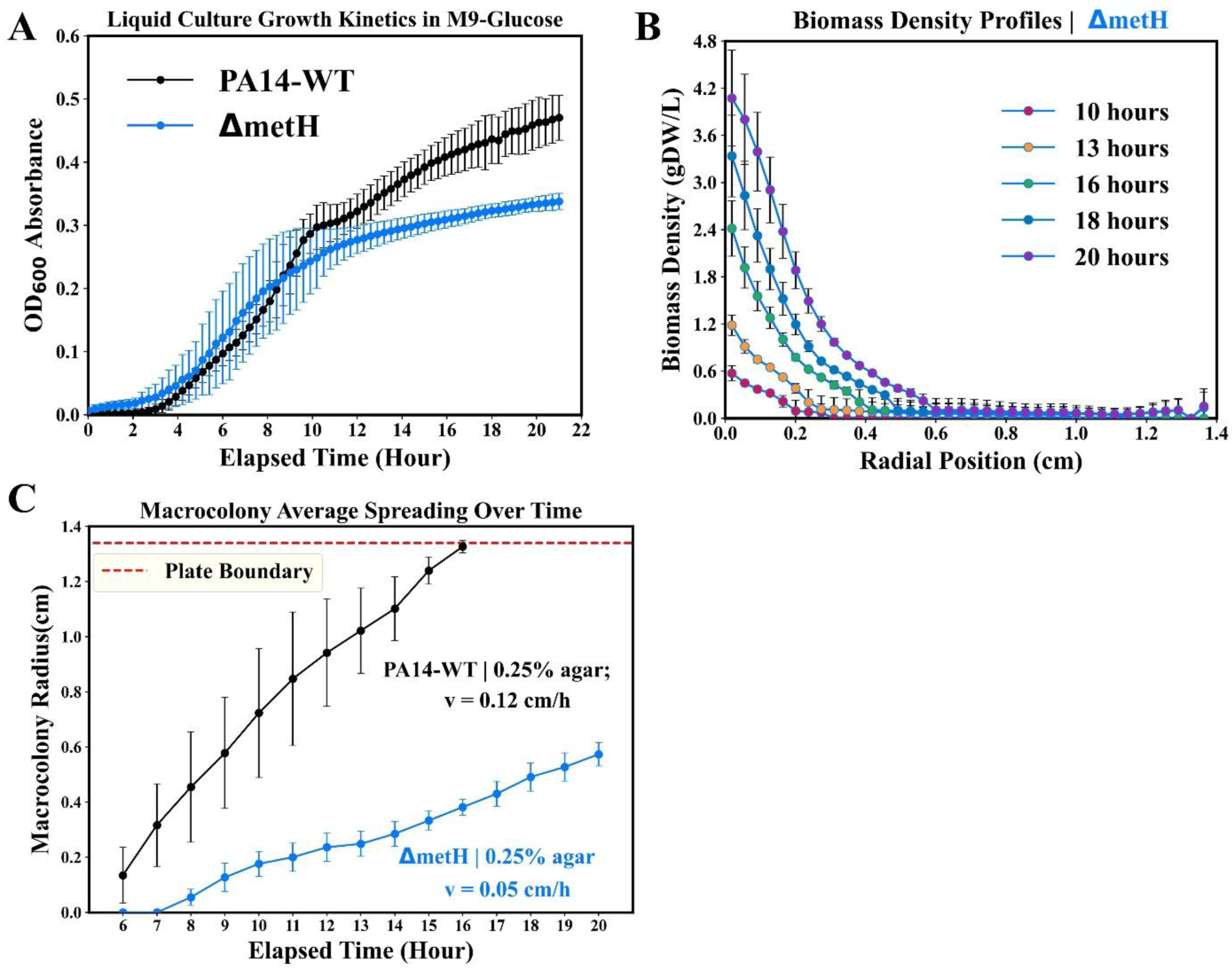
ΔmetH growth curves and timelapse swim plate assay measurements for ΔmetH macrocolonies. (**A**) WT and ΔmetH growth curves assays performed in 96-well reader plates with M9-glucose media. OD_600_ average and standard deviation determined from n = 6 batch replicate wells. (**B**) Average ΔmetH macrocolony density profiles in 0.25% agar swim plate assays measured at specified time points for n = 3 swim plate assays. (**C**) ΔmetH macrocolony average spreading over time compared against WT average macrocolony spreading (values transferred from Fig 4).

The overlapping ΔmetH and WT OD_600_ measurements across most of the expected exponential growth phase supported the speculation that ΔmetH had better growth fitness in M9-glucose than other growth-inhibiting KO groups. ΔmetH growth inhibition did become notable halfway through the assay as ΔmetH OD_600_ plateaued while growth in the WT group continued to increase cell culture opacity. We suspect that exogenous L-methionine in M9-glucose media was temporarily able to supply ΔmetH demands sustain growth near WT levels until L-methionine was depleted in the media. Confined expansion was also confirmed with ΔmetH macrocolonies observed with average spreading velocity approximately two times slower than WT macrocolonies (**Fig 8C**). From the inverse relationship between motility and growth rate in the Fisher-KPP traveling wave speed (**Eq. 2**), the decrease in spreading velocity would require ΔmetH continually experiencing an order of magnitude reduction to growth rate if one assumed that *metH* KO did not affect bacterial motility at the leading macrocolony edge.

Given that cells at the leading edge would have access to nutrients in the uncolonized swim plate and growth dynamics between ΔmetH and WT were similar at the start of liquid cultures, it is unlikely that ΔmetH macrocolony development was solely affected by growth inhibition and suggests *metH* KO inhibited growth and bacterial motility. Connections between *metH* and active bacterial motility have been reported by Yeung *et al.* where gene KOs in the methionine biosynthesis pathway inhibited PA14 motility in low weight agar gels [63]. We hypothesize that the underlying interaction between methionine biosynthesis and bacterial motility is also mediated through SAM. For model flagellated bacteria, SAM supplies methyl groups for methylation reactions between receptor proteins and the enzyme product of the *cheR* motility gene [64]. The dynamic balance in receptor protein methylation states through *cheR*-SAM helps regulate cell signaling to flagellar motor structures controlling flagella switching between counterclockwise and clockwise rotation, and consequently, changing the swimming trajectory for flagellated bacteria [65, 66, 67, 68]. This disruption to swimming motility regulation may make swim agar navigation more difficult for ΔmetH as frequent swimming trajectory changes have been highlighted as critical motile cell behavior for effective mean displacement in confined, porous geometries [69, 70, 71]. Future GEM-CM iterations could leverage SAM metabolite flux from FBA outputs to predict the *D_B_* parameter values, though these efforts will need to be supported with individual cell tracking studies of WT and ΔmetH within gel environments.

## 4. CONCLUSIONS

In this work, we developed and presented GEM-CM, a depiction of spatially heterogenous bacteria populations with metabolic capabilities informed by a coarse-grained GEM. Through its application of RF regression to reduce the complexity involved with coupling GEMs to dynamic PDE environments, GEM-CM can allow leaner PDE systems to incorporate mechanistic metabolic information into their growth rate and metabolite production predictions. The three carbon source inputs selected via feature importance were able to train ensemble models that could effectively inform reaction rate values in the PDE, and through its support during simulation runs, allowed GEM-CM to capture the spatiotemporal expansion of PA14 swim plate macrocolonies over long time scales. We demonstrated GEM-CM applications to probe PCN production behavior in PA14 populations by contextualizing assay observations against model outcomes of different FBA-simulated PCN synthesis. We expect that simulated metabolic activity in the GEM-CM framework can be extended outside of PCN production and potentially represent different GEM metabolic ‘states’, defined via user-imposed constraints or -omics data, that help interrogate heterogeneous cell activity in microbial populations. Lastly, we showed that ensemble coarse-graining was adaptive to FBA solution spaces when *in silico* gene KO contexts were imposed and GEM-CM successfully predicted biomass growth and PCN production inhibition trends in KO macrocolonies. Motility assay screenings further revealed prospective L-methionine metabolic pathways that could disrupt PA14 growth as well as decrease population spreading efficacy in porous, semisolid media.

Continuing work with GEM-CM could seek to widen the scope of interpretable simulation outputs beyond bulk averages and biomass density distribution by employing *in vitro* measurements capable of preserving spatial granularity and monitoring multiple compounds, especially nutrient sources to better frame experiment conditions against simulation predictions. From a methodological perspective, future evaluations of ensemble performance under varying number of feature inputs are also motivated to better enumerate the constraints influencing GEM-CM’s depicted utility. For further investigations with *P. aeruginosa*, we expect that elucidating the intracellular and environmental mechanisms that influence bacterial motility [72, 73, 74] would enhance how GEM-CM simulations depict pathogenic populations in biological settings like obstructive respiratory mucus layers. This direction was briefly explored during our model framework development by altering *D_B_* to reflect lower PA14 motility in 0.28% w/v agar (*Supplementary Materials, Appendix H*), which had GEM-CM producing model density profile predictions with accuracy similar to the presented 0.25% agar case (**Fig S.7**) as well as reductions to simulated biomass growth and PCN production that matched swim plate experiment measurements (**Fig S.8**). Overall, we envision GEM-CM contributing to the current suite of multi-scale, dynamic GEM modeling as an early tool for describing microbial systems with minimal *a priori* information and potentially informing which salient variables to monitor during experimental platform preparation.

## 5. MATERIALS & METHODS

### 5.1: Bacteria strains and growth conditions

WT *P. aeruginosa* PA14 was generously donated by the Goldberg lab [75]. PA14 strains from the non-redundant transposon library developed by Liberati *et al.* [53] utilize a MAR2xT7 transposon element to render their target gene inactive. Each PA14 KO transposon insertion site in the active gene is reported in **Table S.1**. All nutrient media and reagents were sourced from Fisher Scientific unless specified otherwise. Nutrient media was supplemented with either 100 μL/mL carbenicillin (Teknova) or 15 μL/mL gentamicin (Gibco) for selective maintenance of the WT strain and transposon KOs, respectively. When performing bacterial assays, PA14 strains were first streaked on LB-Miller agar plates to raise single colonies and then transferred to LB-Miller broth for overnight incubation at 37°C with 150 RPM agitation (Excella E24, New Brunswick Scientific). Overnight cultures were diluted in fresh LB-Miller broth for PA14 to regain exponential growth phase (∼0.5 OD_600_) before their transfer to swim plate assays.

### 5.2: Swim plate assay preparation

Granulated agar was heated in MilliQ water until agar granules had fully melted and combined with stocks of M9-glucose media components (**Table S.2**). After vortex mixing, the resulting gel solution had desired 0.25% agar weight percentage, appropriate antibiotic supplement, and the nutrient composition of M9-glucose. Swim plate agar gels, approximately two mm in thickness, were formed from three mL of molten gel solution were transferred to 6-well plate dishes (Corning) and set aside for 30 minutes at room temperature to solidify. PA14 WT or transposon KO was transferred from exponential phase LB-Miller liquid cultures to the center of the well dish using a 20-µL micropipette tip to pierce through the vertical length of the gel. Swim plates that were prepared for single time point measurement were kept in a 37°C static incubator (ThermoScientific) until their intended measurement time. Swim plates prepared for timelapse imaging were first placed in the static incubator for 6 hours and then transferred to the widefield microscope.

### 5.3: Swim plate assay imaging and density profile analysis

Detailed procedures for swim plate assay imaging and image processing are provided in *Supplementary Materials*, *Appendix C*. Briefly, tiled brightfield images of well plate dishes were collected over a constant z-position plane with a Leica Thunder Imager Live Cell microscope and Leica K5 CMOS camera. Swim plate assays designated for timelapse imaging were first secured to an open-bottom, temperature-controlled chamber set to 37°C and mounted to the microscope stage. Swim plate development during timelapse experiments was measured between 6 and 20 hours at one hour intervals. Macrocolony intensity profiles were calculated from the tiled brightfield images using the public ImageJ plugin ‘Radial Profile Plot’ setting the center point as the assay’s inoculation point site. Brightfield intensities were converted to gDW/L using a power law relationship determined from a calibration curve (**Fig S.1**). Power law exponent and slope parameters were optimized such the average biomass density from the estimated macrocolony radial profile at t = 20 hours agreed with their respective gDW/L density measurements (covered in following section). We also assumed that power law parameters for intensity-biomass conversion did not fluctuate over time when estimating biomass density profiles at time points earlier than t = 20 hours.

The radial position origin for experimental profiles was adjusted to be approximately one mm after the swim plate inoculation mark due to the lingering cavity formed by the pipette stab. Macrocolony spreading distance was determined to be the farthest position that the biomass density profile was continuously above a 0.1 gDW/L threshold, the lowest density of PA14 distinguishable from agar background under brightfield illumination. Bulk macrocolony biomass densities for time points earlier than t = 20 hours were calculated by dividing the radial biomass density profile’s double integral by well plate area (**Eq. 3**). Spreading velocity in swim agar gels was estimated from timelapse images using a linear best-fit of macrocolony spreading distance over time across n ≥ 5 time points (**Fig 3B**).

### 5.4: Measurement of sample biomass density

Procedures reported in Kim *et al.* were repeated to measure total protein concentrations in the bulk swim agar and approximate the total protein-gDW/L conversion [35]. Swim plate gels were broken up using a 3-mL Leur-Lok syringe (BD Diagnostic) and an additional two mL of Gibco phosphate buffered saline (ThermoFisher) to help lift as much of the sample from the well dish surfaces. Once solids were separated from supernatant via centrifugation (Eppendorf 5425R, 5000g, eight minutes, and 4°C), solid fractions were thoroughly washed with cold PBS and pelleted 3 times. Solids were then resuspended with two mL of lysis buffer [10 mM Tris-HCl, 100 mM NaCl, 1.0 mM EDTA, pH 7.4] and transferred to O-ring tubes (Corning) containing 425-600 μm glass beads (Sigma). Samples were mechanically lysed using a Qiagen Tissuelyser LT homogenizer (five cycles of one-minute duration at 50 Hz) and prepared, in triplicate, for Pierce^TM^ bicinchoninic acid assay (Thermo Fisher) measurements following the manufacturer instructions. Sample 562 nm absorbance quantified using a TECAN Infinite 200 PRO plate reader was converted to total protein concentration and then converted to gDW/L approximating that protein made up half the PA14 dry cell weight [35, 76]. Average gdW/L measurements for each WT and KO group are listed in **Tables S.4 & S.5**.

### 5.5: Measurement of pyocyanin concentration

Decanted supernatant fractions from homogenized swim plate assays were retained and filtered through a 0.22-μm PDVF membrane (NEST Scientific). Adapting PCN isolation procedures from Fong *et al.* [77], two mL of chloroform was added to supernatant samples and vortex mixed to pull aqueous PCN into the organic phase. The aqueous phase was removed and 1.2 mL of 0.1 M HCl was vortexed mixed in with the remaining organic phase. The HCl fraction containing PCN was transferred to a 96-well plate and quantified through 520 nm absorbance measured in triplicate by the TECAN plate reader. A calibration curve of micromolar PCN concentration vs 520 nm absorbance was collected with purified PCN (Sigma, CAS 85-66-5). Average PCN concentrations for each WT and KO group are listed in **Tables S.4 & S.5**.

### 5.6: Partial differential equation system for genome-scale, metabolism-informed continuum model *simulated swim plates*

The PDE in the GEM-CM continuum layer is defined over 1-D cylindrical spatial mesh with the left and right boundaries representing the center and non-permeable well dish wall, respectively. PA14 (*C_B_*) grows with growth rate *v_biom_* and spreads across and the model space with diffusivity *D_B_* (**Eq. 4**). The macrocolony population’s traveling wave properties, see *Supplementary Materials, Appendix F* for further discussion, supported the empirical estimation of *D_B_* values through the Fisher-KPP traveling wave velocity equation and measured macrocolony spreading velocity (Table S.5). PA14 growth is sustained by three carbon sources evaluated to be of high importance to the ensemble model. The diffusion and consumption of each carbon source follow the general form defined in **Eq. 5** with their own unique *v_N,i_* that are defined through concentration conversions into ensemble inputs (**Eq. 1**). Lastly, PCN is generated by the growing PA14 population with a production rate of *v_PCN_* (**Eq. 6**). We assume that metabolite diffusion (*D_N_*) is equal for all non-PA14 species, not altered by biomass or agar presence, and not affected by interactions between metabolites.

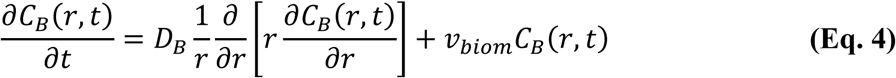

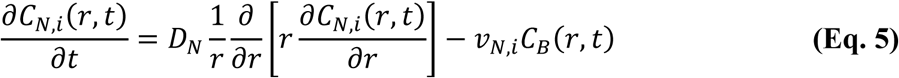

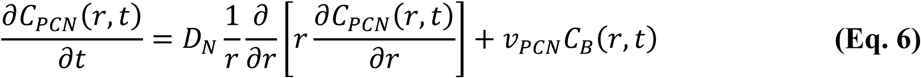

Boundary conditions for all PDE variables had their flux at the center and at the plate boundary (*R*) set to 0 (**Eq. 7**). All nutrients were evenly distributed at their starting M9-glucose concentrations (**Eq. 8, Table S.3**), the initial PA14 inoculum in the simulation was described using a step function (**Eq. 9**), and PCN was not initially present in the model space at the start of simulation (**Eq. 10)**. Reaction rate values *v_biom_*, *v_N,i_*, and *v_PCN_* were updated by the ensemble model at five minute intervals using the discrete concentrations of each carbon source, and GEM-CM simulations could only use one assigned ensemble throughout a batch run. The PDE system is solved using the finite volume solver package FiPy (v 3.4.2.1) [78]. Additional details regarding spatial and time discretizations are provided in *Supplementary Materials*, *Appendix F* and **Table S.8**.

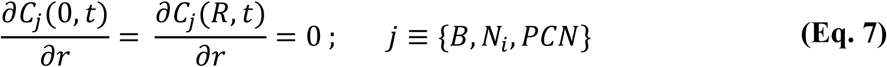

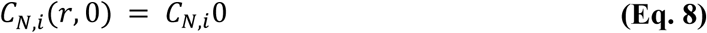

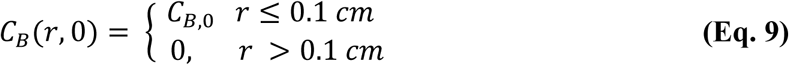

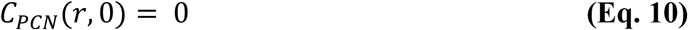

### 5.7: FBA simulation and ensemble training in the GEM-CM coarse-grain layer

The PA14 GEM was generated from the genome-scale metabolic network reconstruction iPau21 using the Cobrapy package (v0.29.0) [79]. PA14 PCN production was interpreted through an added demand reaction operating as a sink drawing all synthesized PCN out of the GEM cytosol compartment. The upper bound for biomass flux was also constrained to 0.9 (i.e. the maximum growth rate set for PA14 was 0.9 hour^-1^); however, no alterations were made to the iPau21 biomass objective function used to evaluate biomass flux. Datasets of FBA simulations were generated by randomly sampling different flux constraints for exchange reactions transporting major carbon source metabolites in M9-glucose growth media [80] (**Table S.6**). PCN synthesis flux inclusion in the FBA objective was determined through a decision tree with rules tailored to permit PCN flux when biomass flux values were between defined minimum and maximum thresholds.

Two dataset collections were compiled for each KO and Γ-rule value condition for different RF regression outputs evaluated using the RandomForestRegressor class in scikit-learn (v1.3.0). The ‘low growth’ FBA simulation dataset with biomass flux ≤ 0.3 hour^-1^ was used to identify which three exchange reactions contributed high importance during regression (**Table S.7**). The ‘high growth’ FBA simulation dataset with biomass flux ≤ 0.9 hour^-1^ was used to train the ensemble model(s) predicting *v_biom_*, *v_N,i_*, and *v_PCN_* reaction rates in GEM-CM simulations. During ensemble training on ‘high growth’ datasets, inputs were the three carbon source exchange reactions with the highest RF feature importance and prediction targets were biomass and PCN fluxes. Additional coarse-graining procedure details and regression validation are discussed in *Supplementary Materials, Appendix E*.

### 5.8: Tuning parameters converting metabolite concentrations to ensemble inputs

Both carbon source consumption rates in the continuum layer and the apparent metabolite flux inputs for the ensemble relied on appropriate *v_max_* and *k_S_* parameter values to define each carbon source’s relationship between concentration and mass flux. To help fit these parameter values, dynamic simulations of the isolated ensemble were conducted to extract carbon source consumption over time predicted by the *v_max_* and *k_S_* set. Predicted metabolite concentrations at discrete time points were then compared to liquid culture data collected by McGill *et al.* [43] in order to evaluate a loss function sum of each carbon source’s root mean squared error normalized to their mean concentration. This loss function was then minimized using Bayesian optimization procedures and constraining the parameter search spaces to 0.5 – 5 mmol/gDW/hour and 1 – 50 µM for *ν_max_* and *k_S_*, respectively, with the exception of glucose *ν_max_* spanning 0.5 – 10 mmol/gDW/hour. More details around the Bayesian operation process are included in *Supplementary Materials, Appendix F.* Final *ν_max_* and *k_S_* parameter values were evaluated as the geometric mean of the top ten candidates with the lowest loss function score (**Table S.9**).

### 5.9: Statistical analysis

Significance between swim plate assays groups conditions were evaluated using 2-sample t-tests with equal variance assumptions confirmed by Levene’s test performed in Python. To avoid favorably biasing model prediction accuracy, coefficient of determination scores between GEM-CM and experiments excluded residual error for data points if experimental quantities were near 0 (i.e. empty swim plate regions of *C_B_* = 0 or depleted metabolite concentration).

## ACKNOWLEDGEMENTS

The authors acknowledge the Rivanna High Performance Computing system for facilitating a high-throughput workflow for GEM-CM model simulations. The authors also gratefully acknowledge: Dr. Mohammad Mazharul Islam for discussions troubleshooting the GEM iPau21; Prof. Shayn Peirce for providing access to microscope equipment resources; and lastly Jenny Vu for data collection and analysis assistance during transposon mutant screening.

## SUPPORTING INFORMATION CAPTIONS

**Fig S.1. Log-log plot of total protein concentration against average brightfield intensity.** Total protein measurements were conducted in triplicate for each PA14-agar sample.

**Fig S.2. Parity plots of ensemble biomass objective function (biomass flux) predictions for hold-out FBA testing data.** Ensemble model results are visualized for (A) WT, Γ = 0.2; (B) WT, Γ = 0.9; (C) ΔphzM; (**D**) ΔargG, and (**E**) ΔmetH. All ΔpurH FBA testing data and ΔpurH ensemble predictions monotonically yielded non-viable PA14 biomass flux as expected.

**Fig S.3. Ensemble regression interpolation of PCN synthesis outputs from iPau21 GEM**. Hold-out FBA testing data (empty markers) are plotted by their biomass objective function and PCN synthesis fluxes in order to illustrate the step function-like behavior produced by the objective function decision tree (Fig 2A). Ensemble predictions (filled markers) for Γ = 0.2 (**A**) and Γ = 0.9 (**B**) testing data illustrate random forest regression outputs that interpolated the different step function behavior.

**Fig S.4. Ensemble coarse-graining of ΔphzM, and ΔpurH metabolic behaviors versus WT**. Heatmaps of biomass objective function and PCN flux predictions from (**A**) WT (Γ = 0.2), (**B**) ΔphzM, and (**C**) ΔpurH ensembles. Glucose and L-glutamate inputs to each ensemble were varied while maintaining L-arginine input flux at −1.4 mmol/gDW/hour.

**Fig S.5. Ensemble coarse-graining of ΔargG metabolic behavior versus WT**. Heatmaps of biomass objective function and PCN flux predictions from (**A**, **B**) WT (Γ = 0.2) and (**C, D**) ΔargG ensembles at varying levels of L-arginine input flux. Glucose and L-glutamate inputs to each ensemble were varied with L-arginine input flux either set to −0.1 mmol/gDW/hour (LOW L-Arg) or −1.4 mmol/gDW/hour (HIGH L-Arg).

**Fig S.6. Ensemble coarse-graining of ΔmetH metabolic behavior versus non-mutant case.** Heatmaps of biomass objective function and PCN flux predictions from **(A, B) non-mutant** and **(C, D)** ΔmetH ensembles at varying levels of L-methionine input flux. The non-mutant ensemble was trained using an FBA dataset without any in silico gene deletion contexts and had glucose, L-arginine, and L-methionine as input features. Glucose and L-arginine inputs to ΔmetH and non-mutant ensembles were varied with L-methionine input flux either set to −0.04 mmol/gDW/hour (LOW L-Met) or −0.8 mmol/gDW/hour (HIGH L-Met).

**Fig S.7. PA14 swim plate assays and simulations in 0.28% agar. (A)** Tiled brightfield images of PA14 macrocolonies (opaque white regions) in 0.28% versus 0.25% swim agar at end of timelapse observation. Scale bar = 2 cm. **(B)** Average macrocolony radii versus time and mean spreading velocity estimations of 0.25% (transferred from Fig 3, black dots) and 0.28% (n = 5, blue diamond) agar swim plates; error bars = 1 standard deviation. Red dashed line denoting the distance to the impermeable well dish wall. **(C - E)** Experimental and GEM-CM predicted biomass density profiles at discrete time points scaled to each group’s average endpoint biomass density. GEM-CM density profile predictions are adjusted by **δ** (two hours). Group average and sample replicate density profile visualization follow the schemes from Fig 3.

**Fig S.8. Effects of reduced PA14 motility on biomass growth and PCN production in GEM-CM predictions**. (**A**) 2-D heatmaps of predicted growth rate over space and simulated time for a 0.25% agar swim plate simulation (*D_B_* = 1.5 E-06 cm^2^/s) and a 0.28% agar swim plate simulation (*D_B_* = 0.5 E-06 cm^2^/s). Cell values denote the average growth rate within a specified radial position range and time point. Radial position increases left-to-right along the heat map and simulated time increases going from top-to-bottom. (**B**) 2-D heat maps for PCN production rates from the same GEM-CM simulation run as their growth rate counterpart in **A**. (**C**) Biomass density and PCN concentrations from 0.28% agar experiments and simulations relative to their mean quantities from the 0.25% agar condition. Scatter plot markers overlaying bar graphs represent each experiment or GEM-CM replicate.

**Table S.1. Mar2xT7 single insertion transposons used in investigation.** PA14 mutant library preparation and validation reported by Liberati, N. T. et al. (2006) Proc. Natl. Acad. Sci. Additional information for the transposon library is provided at https://pa14.mgh.harvard.edu/cgi-bin/pa14/home.cgi.

**Table S.2. Preparation of M9-glucose media from stock solutions.** See *Supplementary Materials, Appendix B* for stock solution preparation procedures.

**Table S.3. Estimated starting carbon source concentrations for M9-glucose media**. Amino acid concentrations estimated from weight percentage compositions reported for casamino acids derived from casein using acid hydrolysis [80].**Table S.4. Bulk quantities for wild type and transposon mutant swim plate assays following single timepoint observation procedures. ^(a)^**Assay measurements (n = 5) collected after 20-hour incubation in 0.25% agar swim plates. Two sample t-tests comparing sample measurements against WT (t = 20 hour) group. **: p < 0.01, ***: p < 0.001. **^(b)^**Assay measurements (n = 3) from 0.25% agar swim plates collected at their reported timepoint post-inoculation.

**Table S.5. Bulk measurements and spreading velocities of swim plate macrocolonies prepared for timelapse observation procedures.** All biomass density and PCN concentrations collected at t = 20 hours. **^(a)^**Two sample t-test performed comparing sample measurements against WT, 0.25% agar group. **: p < 0.01, ***: p < 0.001.

**Table S.6. FBA simulation constraints sampled for M9-glucose growth media.** See *Supplementary Materials, Appendix E* for random sampling discussion.

**Table S.7. Ensemble model parameters and their corresponding metabolites for GEM-CM simulations. ^(a)^**Regression criterion applied Friedman mean squared error in place of standard squared error. **^(b)^**Monotonic, non-viable FBA training data had ensemble parameters default to lowest sample numbers for leaf node and decision tree splitting. Input features and GEM-CM metabolites were defaulted to repeating WT inputs.

**Table S.8. GEM-CM continuum layer variables for simulation mesh discretization and timesteps.**

**Table S.9. Diffusion coefficients for GEM-CM continuum layer. ^(a)^**Bacteria motility coefficients identified for each swim plate assay replicate using its measured macrocolony spreading velocity. A GEM-CM batch simulation was performed for each bacteria diffusivity value in a gel composition group with all other model parameters and initial conditions held constant.

**Table S.10. Monod kinetics parameters for coupling ensemble and continuum layers in GEM-CM.** Values are the geometric means of the top ten candidates from Bayesian optimization procedures (*Supplementary Materials, Appendix F*).

**Table S.11. Flux variability analysis of select reactions in the iPau21 GEM.** See *Supplementary Materials, Appendix G* for flux variability analysis discussion.

## Notes

https://github.com/morganskim2/GEMCM_2026

https://doi.org/10.5281/zenodo.23897610

### Competing Interest Statement

The authors have declared no competing interest.

### Summary of Updates

Supplemental materials have been uploaded.

