## Supplementary material for "Coarse-graining genome-scale metabolic models for dynamic, multi-scale simulations of *Pseudomonas aeruginosa* in gel environments": Supplmentary Materials_001

### ***Appendix A****: Single transposon insertion PA14 mutants from prepared, non-redundant library* [1]*.*

**Table S.1. Mar2xT7 single insertion transposons used in investigation***.*

| **Mutant ID in Library** | **Active Gene Name** | **Hypothesized Insertion Position within Active Gene (bp) /**  **Active Gene Size (bp)** |
| --- | --- | --- |
| 26056 | *argG* | 1240 / 2325 |
| 39111 | *metH* | 67/ 3705 |
| 40343 | *phzM* | 239 / 1005 |
| 46986 | *purH* | 129 / 1608 |

PA14 mutant library preparation and validation reported by Liberati, N. T. *et al.* (2006) *Proc. Natl. Acad. Sci.* Additional information for the transposon library is provided at https://pa14.mgh.harvard.edu/cgi-bin/pa14/home.cgi.

### ***Appendix B****: Defined minimal media for swim plate assay experiment.*

M9-minimal salts (5x, Sigma) were dissolved and autoclaved in MilliQ water according to manufacturer specifications. Separate stock magnesium sulfate (1 M, Sigma), calcium chloride (0.1 M, Sigma), D-glucose (20% w/v, Sigma), and iron (II) sulfate heptahydrate (0.1 M, Fisher) solutions were also prepared with MilliQ water and filtered through a 0.22-µm PVDF membrane. We additionally included casamino acids (Fisher) solution in the growth media to further supplement PA14 growth. Batches of M9-glucose growth media were prepared from these stock solutions the day of swim plate assays following the relative dilution scheme listed in **Table S.2**. Starting concentrations of carbon sources (**Table S.3**) were estimated using weight percentage compositions reported for casamino acids derived from casein using acid hydrolysis [2]. To achieve a molten state with M9-glucose agar, granulated agar was first heated in MilliQ water alone and all M9-glucose stock solutions were mixed into the high weight percentage agar solution once the molten agar was cool enough to handle.

**Table S.2**. **Preparation of M9-glucose media from stock solutions.**

| **Stock Solution** | **Dilution for**  **M9-Glucose-Gent** | **Dilution for**  **M9-Glucose-Carb** |
| --- | --- | --- |
| M9-minimal salts (5x) | 1:5 | 1:5 |
| MgSO_4_ (1 M) | 1:1000 | 1:1000 |
| CaCl_2_ (0.1 M) | 1:1000 | 1:1000 |
| FeSO_4_ ∙7H_2_O (0.1 M) | 1:1000 | 1:1000 |
| Glucose (20% w/v) | 1:50 | 1:50 |
| Casamino acids (1% w/v) | 1:10 | 1:10 |
| Carbenicillin (100 mg/mL) |  | 1:1000 |
| Gentamicin (50 mg/mL) | 1:3000 |  |

**Table S.3. Estimated starting carbon source concentrations for M9-glucose media***.*

| **Metabolite** | **Concentration (mM)** |
| --- | --- |
| Glucose | 22.20 |
| L-Alanine | 0.67 |
| L-Arginine | 0.25 |
| L-Asparagine | 0.57 |
| L-Aspartic acid | 0.56 |
| L-Glutamate | 1.70 |
| L-Glutamine | 1.71 |
| L-Glycine | 0.51 |
| L-Histidine | 0.28 |
| L-Isoleucine | 0.70 |
| L-Leucine | 1.10 |
| L-Lysine | 0.93 |
| L-Methionine | 0.32 |
| L-Phenylalanine | 0.38 |
| L-Proline | 1.67 |
| L-Serine | 1.07 |
| L-Threonine | 0.74 |
| L-Tyrosine | 0.36 |
| L-Valine | 0.94 |

### ***Appendix C****: Swim plate imaging and image data conversion to biomass density.*

Swim plate assays for each PA14 test group were carried out as batch replicates on one 6-well plate. Assays were imaged using a Thunder Imager Live Cell microscope (Leica, 4x/0.10 NA) as either a single timepoint immediately before their downstream sample processing or as a timelapse observation collecting images at one hour intervals between 6-20 hours post-gel inoculation. During timelapse observations, the 6-well plate was secured inside a temperature-controlled, open-bottom chamber with an overhead glass lid; the chamber itself was maintained at 37**°C** and supplied with humid air by a connected air pump. For single timepoint collections, the 6-well plate(s) remained in a stationary incubator set to 37**°C** for the assay duration until the desired endpoint had been reached. Tiled images were then collected with the objective lens kept a constant z-position (~1.2 mm from the well plate bottom) and combined over the x-y space using the instrument program’s ‘Mosaic Merge’ tool. Merged images were cropped in ImageJ such that the well plate dish was an inscribed circle framed by the borders of the ~3.6 cm x ~3.6 cm square image. These image files can be accessed through this investigation’s GitHub repository.

Brightfield intensity in swim plate images had an apparent inverse relationship to local cell density. To establish a calibration curve between intensity and cell density, we prepared well-mixed PA14-agar samples using 30-mL liquid PA14 cultures resuspended to varying cell densities in M9-glucose agar (0.25% w/v) on 12-well dish plate (Corning). Image collection replicated microscopy procedures and instrument settings used for swim plate assay experiments, and brightfield intensity was quantified over a region of 1.1 cm x 1.1 cm positioned near the well dish center. Total protein concentration was measured for each PA14-agar sample following procedures described in *Materials and Methods*. **Fig S.1** visualizes the log-log plot of total protein and average intensity over the 1.1 cm x 1.1 cm region.

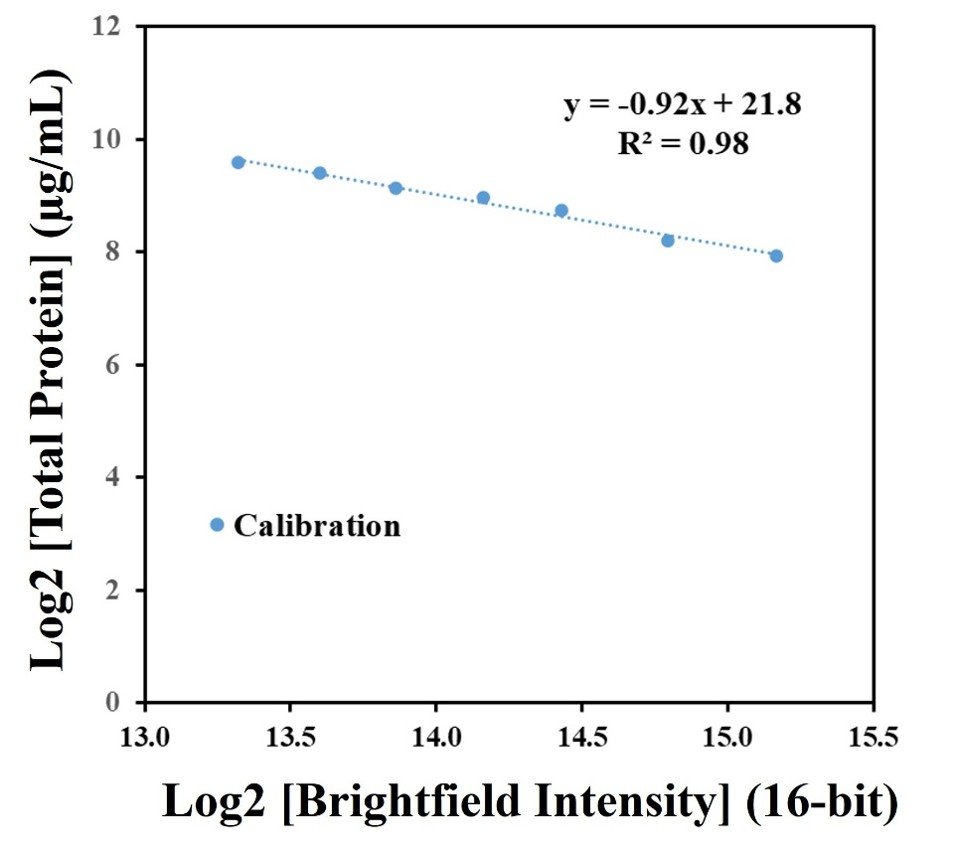

**Fig S.1. Log-log plot of total protein concentration against average brightfield intensity.** Total protein measurements were conducted in triplicate for each PA14-agar sample.

The log-log plot of total protein concentration (*C_prot_*) and brightfield intensity (*I*) was determined to have a linear relationship, and thus a power law function (**Eq. S1**) was applied to estimate total protein concentration from 16-bit brightfield intensity. We attempted to concentrate cell densities in PA14-agar samples to better calibrate the low intensity regime measured in swim plate images, but we were not successful. Thus, due to concerns of high extrapolation error for total protein concentration at these low intensity values, we calculated separate power law parameter values for each swim plate replicate using a minimization objective (**Eq. S2**).

$$C_{prot}(I)=B{(I)}^{A}+\varepsilon$$

**(Eq. S1)**

$$\min_{} \left\{ C_{prot, exper}-\frac{\int_{0}^{R} \int_{0}^{2\pi} \left[ C_{prot, sample}\left( r \right)-C_{prot, bkgrd}\left( r \right) \right] r d\theta dr}{\pi R^{2}} \right\}$$

**(Eq. S2)**

Through **Eq. S2**, the parameters *A* and *B* would be optimized such that the background-subtracted *C_prot_* profile for a swim plate assay image at 20 hours would yield an integrated bulk protein concentration agreeing with that replicate’s total protein measurement (*C_prot,exper_*). Informed by linear regression results from **Fig S.1**, domains for the power law exponent (*A*) and slope (*B*) were constrained to (-1, 0) and (0,1e5), respectively. The error term (ε) was set as the calibration curve fitting error and used as a supporting constraint for *A* and *B* values in order to reinforce near-zero protein concentration at 16-bit intensity saturation. For image data from timelapse assay collections, we assumed that the power-law relationship did not change over time and we uniformly applied the optimized *A* and *B* parameters to estimate all protein concentration profiles in a swim plate replicate’s collection.

To define brightfield intensity at discrete radial positions, we applied the ImageJ plugin ‘Radial Plot Profile’ to calculate intensity profiles(s). The assay inoculation point, visually determined using the lingering gel mark left by the transfer pipette, was set to be the plugin macro’s x-y coordinates for the center point. Background intensity profiles were **determined using a merged image of an uninoculated swim plate collected with identical** microscopy procedures and instrument settings as swim plate assay experiments. The background image was then oriented and **cropped to the same pixel dimensions as macrocolony images in order to repeat plugin macro x-y coordinates and acquire background intensity profiles with spatial contexts comparable to macrocolony profiles.** Prior to parameter optimization, both background and macrocolony profiles had their intensity values averaged over ~360 µm intervals. Once optimal power-law parameters were identified, the **transformed** total protein concentration profile was converted to grams dry weight per liter (gDW/L) estimating that protein comprised half the cell dry mass [3, 4]. When comparing experiment profiles to model predictions in this study, swim plate biomass density distributions were scaled to their respective replicate’s endpoint bulk biomass density; nonetheless, experiment profiles with their gDW/L total values (as depicted in **Fig 5**) can be accessed through the GitHub repository.

### ***Appendix D****: Swim plate assay sample measurements.*

Representative bulk biomass density and pyocyanin (PCN) concentration for each swim plate assay replicate were estimated as the average of triplicate measurements. **Table S.4** and **Table S.5** report group averages of single timepoint and timelapse experiments, respectively. When comparing relative differences between swim plate groups, bulk biomass density and PCN concentrations were scaled to the PA14 wild type (WT) 0.25% agar group averages collected under single or timelapse observation protocols. Biomass density profiles, on the other hand, were scaled by their respective replicate’s endpoint bulk biomass measurement prior to group averaging. Additionally, macrocolony spreading velocity for each assay was calculated from timelapse experiments for later estimation of bacterial motility parameters.

**Table S.4. Bulk quantities for wild type and transposon mutant swim plate assays following single timepoint observation procedures.**

| **Sample Group** | **Biomass Density**  (gDW/L) | **Pyocyanin**  (µM) |
| --- | --- | --- |
| WT ^(a)^  (t = 20 hour) | 1.27 ± 0.14 | 47.8 ± 15.0 |
| ΔphzM ^(a)^ | 1.34 ± 0.11 | 1.3 ± 0.2  (**) |
| ΔargG ^(a)^ | 0.04 ± 0.01  (***) | 1.4 ± 0.2 E-02  (**) |
| ΔmetH ^(a)^ | 0.32 ± 0.04  (***) | 5.8 ± 0.7 E-02  (**) |
| ΔpurH ^(a)^ | 0.02 ± 0.01  (***) | 1.1 ± 0.1 E-02  (**) |
| WT ^(b)^  (t = 16 hour) | 0.50 ± 0.02 | 22.2 ± 2.0 |
| WT ^(b)^  (t = 10 hour) | 0.09 ± 0.01 | 2.6 ± 0.1 |

(**a**) Assay measurements (n = 5) collected after 20-hour incubation in 0.25% agar swim plates. Two sample t-tests comparing sample measurements against wild type (t = 20 hour) group. **: *p* < 0.01, ***: *p* < 0.001.

(**b**) Assay measurements (n = 3) from 0.25% agar swim plates collected at their reported timepoint post-inoculation.

**Table S.5. Bulk measurements and spreading velocities of swim plate macrocolonies prepared for timelapse observation procedures.**

| **Sample Group** | **Biomass Density**  **(gDW/L)** | **Pyocyanin**  **(µM)** | **Spreading Velocity (mm/hour)** |
| --- | --- | --- | --- |
| Wild type  (0.25% agar, n = 5) | 1.16 ± 0.21 | 11.6 ± 2.6 | 1.2 ± 0.2 |
| Wild type  (0.28% agar, n = 5) | 0.54 ± 0.07  (**) | 5.9 ± 1.2  (**) | 0.7 ± 0.1  (**) |
| ΔmetH  (0.25% agar, n = 3) | 0.28 ± 0.05  (***) | 2.4 ± 0.5  (**) | 0.5 ± 0.2  (**) |

All biomass density and pyocyanin concentrations collected at t = 20 hours.

(**a**) Two sample t-test performed comparing sample measurements against wild type, 0.25% agar group. **: *p* < 0.01, ***: *p* < 0.001.

### ***Appendix E****: Processes in coarse-grain layer of genome-scale, metabolism-informed continuum models*.

***E.1*** *Flux balance analysis.*

The genome-scale metabolic network reconstruction iPau21 [5] was converted to a PA14 genome-scale metabolic model (GEM) using Cobrapy (v0.29.0) [6]. With the exceptions of adding a demand reaction for PCN and setting the biomass flux upper bound to 0.9 hour^-1^, no other alterations were deemed necessary to the GEM due to a prior investigation’s intensive curation and validation efforts for iPau21 [5]. The objective function for flux balance analysis (FBA) was selected through decision tree rules (**Fig 2A**) to either maximize biomass flux or the sum of biomass and PCN fluxes. An M9-glucose growth media context was simulated by assigning non-zero lower boundaries of exchange reactions transporting major metabolites, ions, and dissolved gases in M9-glucose (**Table S.6**). We note that a negative lower bound for exchange reactions represents mass flux entering the GEM from the extracellular media. All other exchange reactions in the GEM had their lower bound set to zero flux conveying that the metabolite(s) cannot enter the GEM from the extracellular media.

**Table S.6. FBA simulation constraints sampled for M9-glucose growth media.**

| **Compound** | **Reaction ID** | **Low Growth Setting**  **(flux lower bound)** | **High Growth Setting**  **(flux lower bound)** |
| --- | --- | --- | --- |
| **H_2_O** | EX_cpd00001_e | -1.0E+3 | -1.0E+3 |
| **H^+^** | EX_cpd00067_e | -1.0E-4 | -1.0E-4 |
| **CO_2_** | EX_cpd00011_e | -1.3E-4 | -1.3E-4 |
| **O_2_** | EX_cpd00007_e | -2.0E-2 | -2.0E-2 |
| **NH_3_** | EX_cpd00013_e | -1.8E+1 | -1.8E+1 |
| **Na^+^** | EX_cpd00971_e | -1.2E+2 | -1.2E+2 |
| **K^+^** | EX_cpd00205_e | -2.2E+1 | -2.2E+1 |
| **Fe^2+^** | EX_cpd00021_e | -1.0E-1 | -1.0E-1 |
| **Ca^2+^** | EX_cpd00063_e | -1.0E-1 | -1.0E-1 |
| **Mg^2+^** | EX_cpd00254_e | -1.0 | -1.0 |
| **Cl^-^** | EX_cpd00099_e | -2.7E+1 | -2.7E+1 |
| **HPO_4_^2-^** | EX_cpd00009_e | -7.9E+1 | -7.9E+1 |
| **SO_4_^2-^** | EX_cpd00048_e | -1.1 | -1.1 |
| **Glucose** | EX_cpd00027_e | rand(-10,0) | rand(-10,0) |
| **L-Alanine** | EX_cpd00035_e | rand(-1.25, 0) | rand(-5, 0) |
| **L-Arginine** | EX_cpd00051_e | rand(-1.25, 0) | rand(-5, 0) |
| **L-Asparagine** | EX_cpd00132_e | rand(-1.25, 0) | rand(-5, 0) |
| **L-Aspartic acid** | EX_cpd00041_e | rand(-1.25, 0) | rand(-5, 0) |
| **L-Glutamic acid** | EX_cpd00023_e | rand(-1.25, 0) | rand(-5, 0) |
| **L-Glutamine** | EX_cpd00053_e | rand(-1.25, 0) | rand(-5, 0) |
| **L-Glycine** | EX_cpd00033_e | rand(-1.25, 0) | rand(-5, 0) |
| **L-Histidine** | EX_cpd00119_e | rand(-1.25, 0) | rand(-5, 0) |
| **L-Isoleucine** | EX_cpd00322_e | rand(-1.25, 0) | rand(-5, 0) |
| **L-Leucine** | EX_cpd00107_e | rand(-1.25, 0) | rand(-5, 0) |
| **L-Lysine** | EX_cpd00039_e | rand(-1.25, 0) | rand(-5, 0) |
| **L-Methionine** | EX_cpd00060_e | rand(-1.25, 0) | rand(-5, 0) |
| **L-Phenylalanine** | EX_cpd00066_e | rand(-1.25, 0) | rand(-5, 0) |
| **L-Proline** | EX_cpd00129_e | rand(-1.25, 0) | rand(-5, 0) |
| **L-Serine** | EX_cpd00054_e | rand(-1.25, 0) | rand(-5, 0) |
| **L-Threonine** | EX_cpd00161_e | rand(-1.25, 0) | rand(-5, 0) |
| **L-Tyrosine** | EX_cpd00069_e | rand(-1.25, 0) | rand(-5, 0) |
| **L-Valine** | EX_cpd00156_e | rand(-1.25, 0) | rand(-5, 0) |

FBA was repeatedly performed with different lower bound constraints for glucose and amino acid exchange reactions in order to build training datasets depicting the FBA solution spaces of PA14 biomass and PCN fluxes for random forest regression. Lower bounds for ion and dissolved gas exchange reactions were not varied as we assumed that carbon source abundance was the major limiting factor for PA14 growth. A Latin hypercube algorithm provided by the scikit-learn (v1.3.0) package was used to randomly sample mmol/gDW/hour constraint values between the numerical ranges listed in **Table S.6**. A portion of hypercube sampling was dedicated to having higher glucose flux entering the GEM, (-10, -5) mmol/gDW/hour, relative to each individual amino acid flux,
(-1,0) mmol/gDW/hour, due to reports that *P. aeruginosa* preferentially consumes amino acids before consuming glucose [7, 8]. Hypercube sampling was also conducted in two batches, annotated as ‘low growth’ and ‘high growth’ settings in **Table S.6**, to assemble exchange reaction constraint values with distinct FBA biomass flux outputs that were leveraged for separate random forest regression applications (*Appendix E.2*). The ‘low growth’ constraints (n = 4000) broadly held biomass flux to ≤ 0.3 hour^-1^ while the ‘high growth’ constraints (n = 6000) extended the range of FBA-simulated biomass flux to 0 ≤ *v_biom_* ≤ 0.9 hour^-1^. These sampled constraint values were inputs to the GEM to generate biomass and PCN flux FBA solution spaces after *in silico* single gene deletions or rule changes to the FBA objective decision tree.

***E.2****: Random forest regression.*

Random forest (RF) regression was performed with the RandomForestRegressor class in the scikit-learn supervised learning module. Using RF regression, we sought to identify the exchange reactions / media metabolites that were important for differentiating when PA14 had viable or non-viable biomass flux and for training a RF regression model, or ensemble, that would inform growth, nutrient consumption, and PCN production during swim plate simulations. To best achieve these goals, the ‘low growth’ FBA dataset(s) were used to determine the 3 salient carbon sources in M9-glucose growth media for predicting growth rate from predicted biomass flux, and the ‘high growth’ dataset(s) were used to prepare the final ensemble model, or ensemble, informing swim plate simulations.

To evaluate which three carbon sources would be used for coarse-graining M9-glucose growth media, all 19 amino acid and glucose exchange reactions were first provided as RF input features for predicting biomass and PCN fluxes. Regression training was repeated for n = 50 stochastic iterations with different decision tree leaf node depth and number of decision trees. After each training iteration, the feature importance of each reaction, calculated during regression(s), was rank-ordered and recorded; the three exchange reactions with the lowest mean rank were interpreted to influence the most salient carbon sources in the growth media and their metabolite was included as a nutrient variable in swim plate simulations. If multiple exchange reactions had comparable average importance rank, the combination of exchanges that could produce better biomass flux prediction accuracy on the training hold-out data was chosen. Once 3 major exchange reactions were identified, the ‘high growth’ FBA dataset was applied to train the final ensemble informing swim plate simulations. In this RF regression, training features were limited to the 3 selected exchange reactions with additional sample weighting for training data points possessing biomass values below the dataset median. RF hyperparameters for decision tree splitting, leaf node size, and number of decision trees were optimized through cross-validated grid-searching method before being exported for use in the ensemble layer of GEM-CM simulations. **Table S.7** summarizes ensemble hyperparameters and final set of input features for different PA14 conditions.

**Table S.7. Ensemble model parameters and their corresponding metabolites for GEM-CM simulations.**

| **Ensemble Model** | **# of Estimators** | **Decision Tree Depth**  **(Minimum *‘n’* in leaf node, Minimum *‘n’* for split)** | **Input Features**  **(GEM Exchange Reactions)** | **GEM-CM**  **Metabolites** |
| --- | --- | --- | --- | --- |
| **WT**  **(Γ = 0.2)** | **500** | **(23, 46)** | **EX_cpd00051_e**  **EX_cpd00023_e**  **EX_cpd00027_e** | **L-arginine,**  **L-glutamate,**  **glucose** |
| **WT**  **(Γ = 0.9)** | **500** | **(6, 43)** | **EX_cpd00051_e**  **EX_cpd00023_e**  **EX_cpd00027_e** | **L-arginine,**  **L-glutamate,**  **glucose** |
| **ΔphzM**  **(Γ = 0.2)** | **500** | **(14, 41)** | **EX_cpd00051_e**  **EX_cpd00023_e**  **EX_cpd00027_e** | **L-arginine,**  **L-glutamate,**  **glucose** |
| **ΔargG**  **(Γ = 0.2)** | **400** | **(5, 34)** | **EX_cpd00051_e**  **EX_cpd00023_e**  **EX_cpd00027_e** | **L-arginine,**  **L-glutamate,**  **glucose** |
| **ΔmetH ^(a)^**  **(Γ = 0.2)** | **450** | **(5, 16)** | **EX_cpd00051_e**  **EX_cpd00060_e**  **EX_cpd00027_e** | **L-arginine,**  **L-methionine, glucose** |
| **ΔpurH ^(b)^**  **(Γ = 0.2)** | **100** | **(5, 11)** | **EX_cpd00051_e**  **EX_cpd00023_e**  **EX_cpd00027_e** | **L-arginine,**  **L-glutamate,**  **glucose** |

(**a**) Regression criterion applied Friedman mean squared error in place of standard squared error.

(**b**) Monotonic, non-viable FBA training data had ensemble parameters default to lowest sample numbers for leaf node and decision tree splitting. Input features and GEM-CM metabolites were defaulted to repeating WT inputs.

With information from three exchange reactions, ensembles were typically able to predict FBA biomass flux with a root mean square error (RMSE) of ~0.1 hour^-1^ (**Fig S.2**). For the scope of this investigation, we elected to constrain the number of training features given that the majority of the random forest feature importance score was delegated to three input variables; however, future studies could attempt to refine RF-FBA training procedures to lower RMSE. When reviewing ensemble predictions for PCN flux, we observed limitations with the RF regression approach as the technique’s innate interpolation efforts lowered prediction accuracy for the binary states of PCN flux in the FBA training data (**Fig S.3**). Nevertheless, despite disagreements between FBA and ensemble behaviors, we believe the differential PCN synthesis predictions across the two Γ settings still illustrated the ensemble’s capacity to express distinct metabolic behaviors that can be applied by genome-scale, metabolism-informed continuum models in order to probe experiment observations. If binary states are regularly encountered in FBA calculations of desired metabolic behavior, one might consider a combination of RF regression and classification models for predicting different targets.

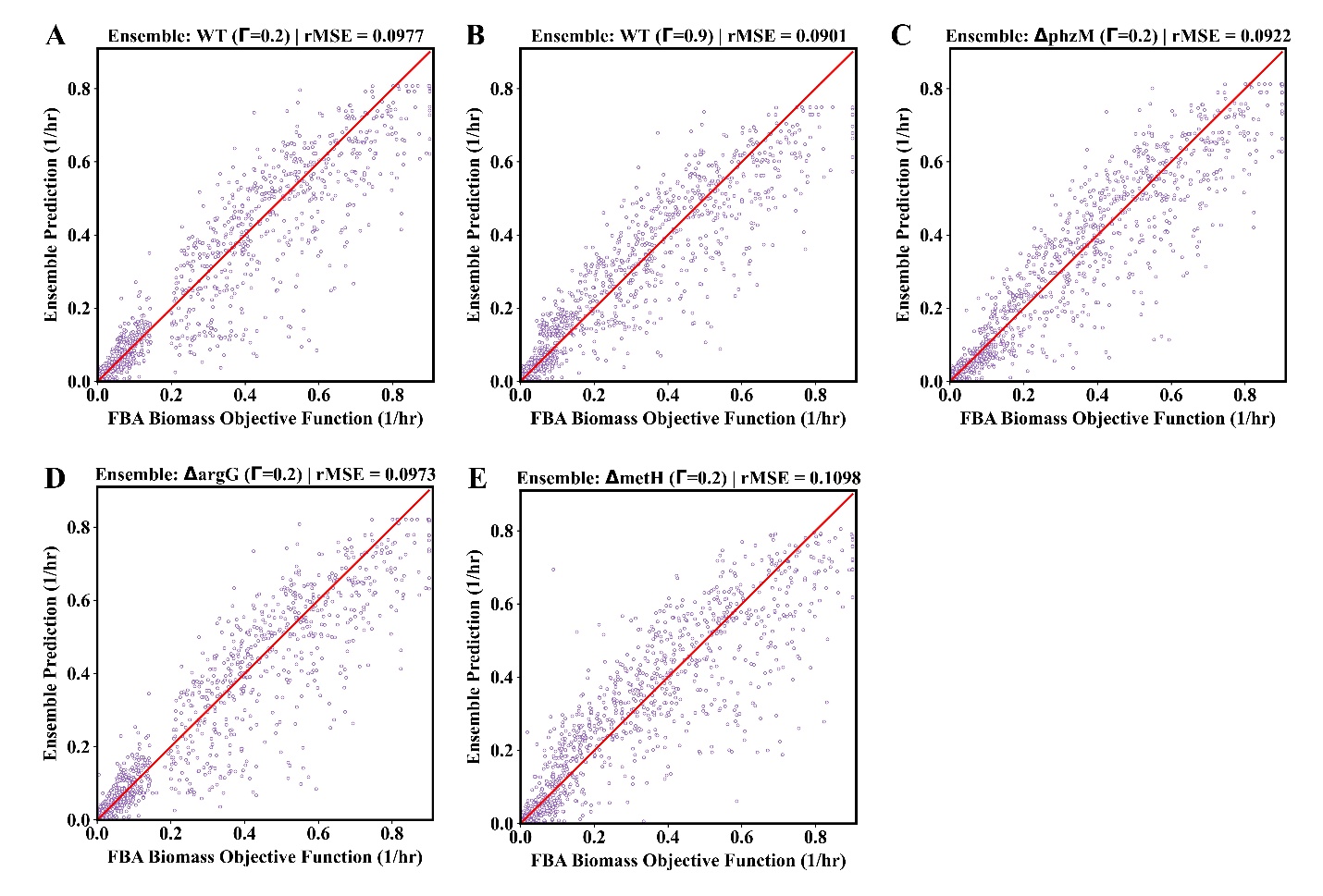

**Fig S.2.** **Parity plots of ensemble biomass objective function (biomass flux) predictions for hold-out FBA testing data**. Ensemble model results are visualized for (**A**) WT, Γ = 0.2; (**B**) WT, Γ = 0.9; (**C**) ΔphzM; (**D**) ΔargG, and (**E**) ΔmetH. All ΔpurH FBA testing data and ΔpurH ensemble predictions monotonically yielded non-viable PA14 biomass flux as expected.

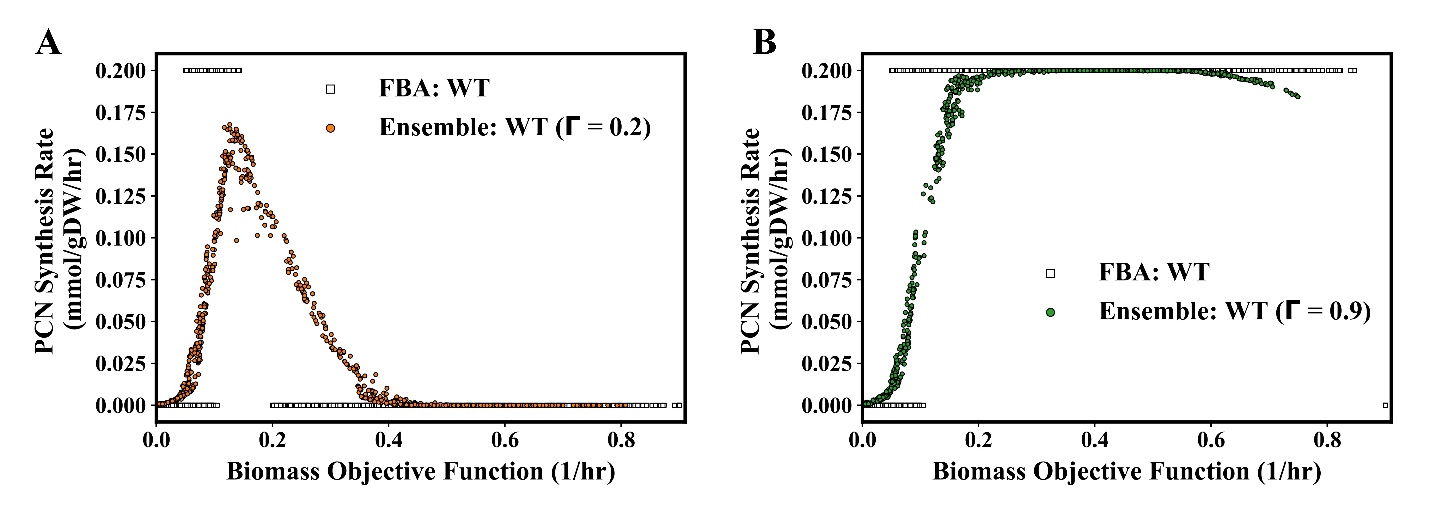

**Fig S.3*.*** **Ensemble regression interpolation of PCN synthesis outputs from iPau21 GEM**. Hold-out FBA testing data (empty markers) are plotted by their biomass objective function and PCN synthesis fluxes in order to illustrate the step function-like behavior produced by the objective function decision tree (Fig 2A). Ensemble predictions (filled markers) for Γ = 0.2 (**A**) and Γ = 0.9 (**B**) testing data illustrate random forest regression outputs that interpolated the different step function behavior.

To confirm that the ensemble models for knockouts had effectively coarse-grained their expected metabolic perturbations, we passed through our regression sets of input values that were designed to extract representative ‘cross-sections’ of biomass and PCN flux predictions from WT and knockout ensembles. These predictions are visualized in **Figs S.4 – S.6** as 2D-binned heatmaps with axes denoting the absolute value of input fluxes. All ensemble PCN predictions were trained on FBA datasets following Γ = 0.2 for objective decision tree rules and usually yielded non-zero prediction values when biomass flux was between 0.1 – 0.4 hour^-1^ (**Fig S.3A**), if PCN synthesis was not perturbed during *in silico* gene deletion FBA. The essentiality of *phzM* to PCN synthesis and *purH* to biomass flux were readily recreated by their knockout ensembles evaluating total PCN or total biomass flux inhibition for all tested inputs (**Fig S.4**). Conversely, the ΔargG ensemble exhibited similar flux predictions to WT when provided with a high L-arginine input (**Figs S.5A & S.5C**); however, ΔargG biomass growth inhibition relative to WT became more evident when ensembles were given lower inputs of L-arginine (**Figs S.5B & S.5D**). Similar prediction variations with amino acid input level were observed with the ΔmetH ensemble (**Figs S.6A & S.6B**) along with notable PCN flux inhibition compared to the non-mutant counterpart. We note that the ‘non-mutant’ ensemble presented in **Fig S.6** was trained using the same base dataset as the WT ensembles in **Figs S.4 & S.5** but the ‘non-mutant’ used glucose, L-arginine, and *L-methionine* as input features to compare predictions against the ΔmetH ensemble. Overall, ensemble regression successfully captured the expected knockout trends from FBA data.

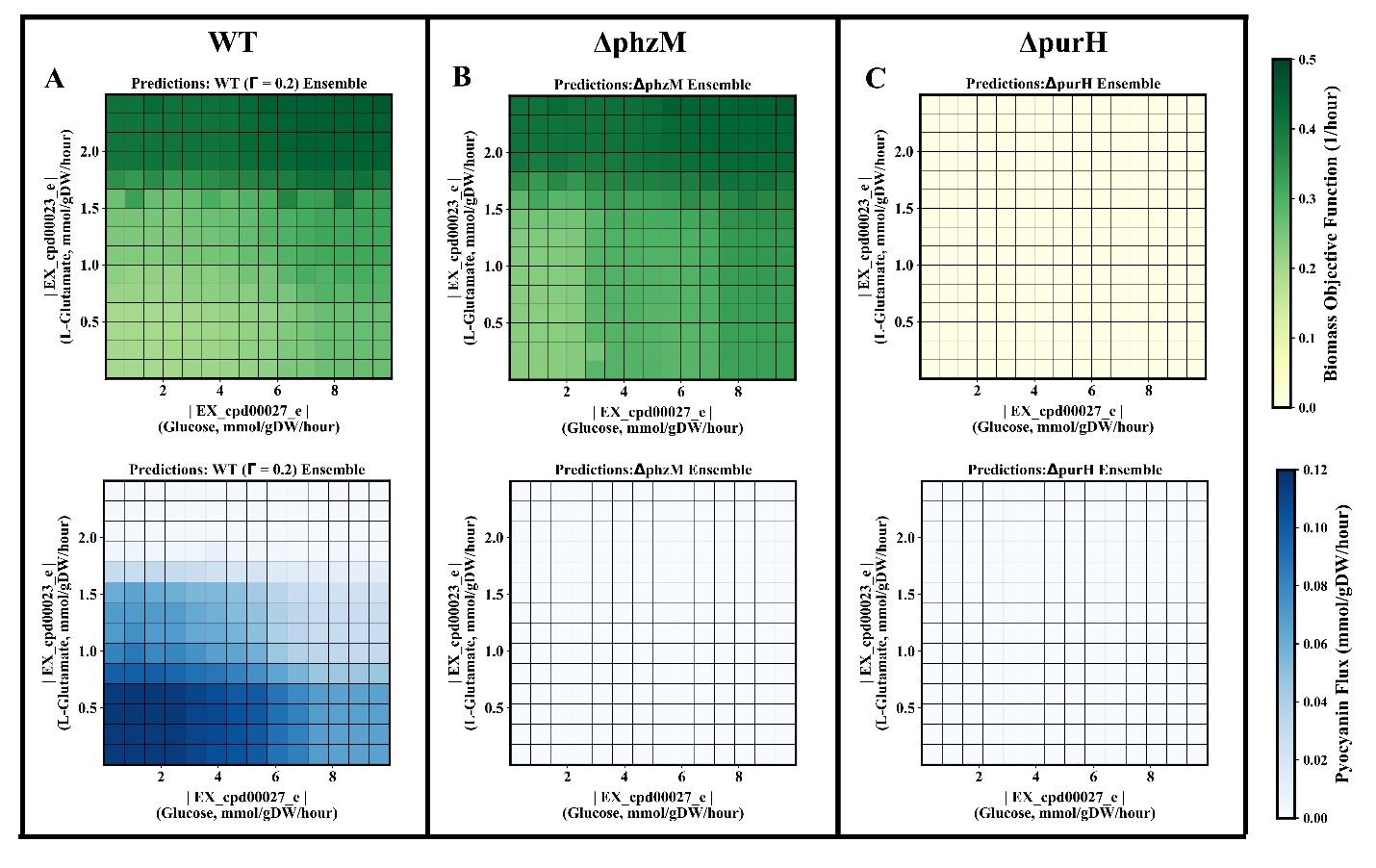

**Fig S.4. Ensemble coarse-graining of ΔphzM, and ΔpurH metabolic behaviors versus WT**. Heatmaps of biomass objective function and PCN flux predictions from (**A**) WT (Γ = 0.2), (**B**) ΔphzM, and (**C**) ΔpurH ensembles. Glucose and L-glutamate inputs to each ensemble were varied while maintaining L-arginine input flux at -1.4 mmol/gDW/hour.

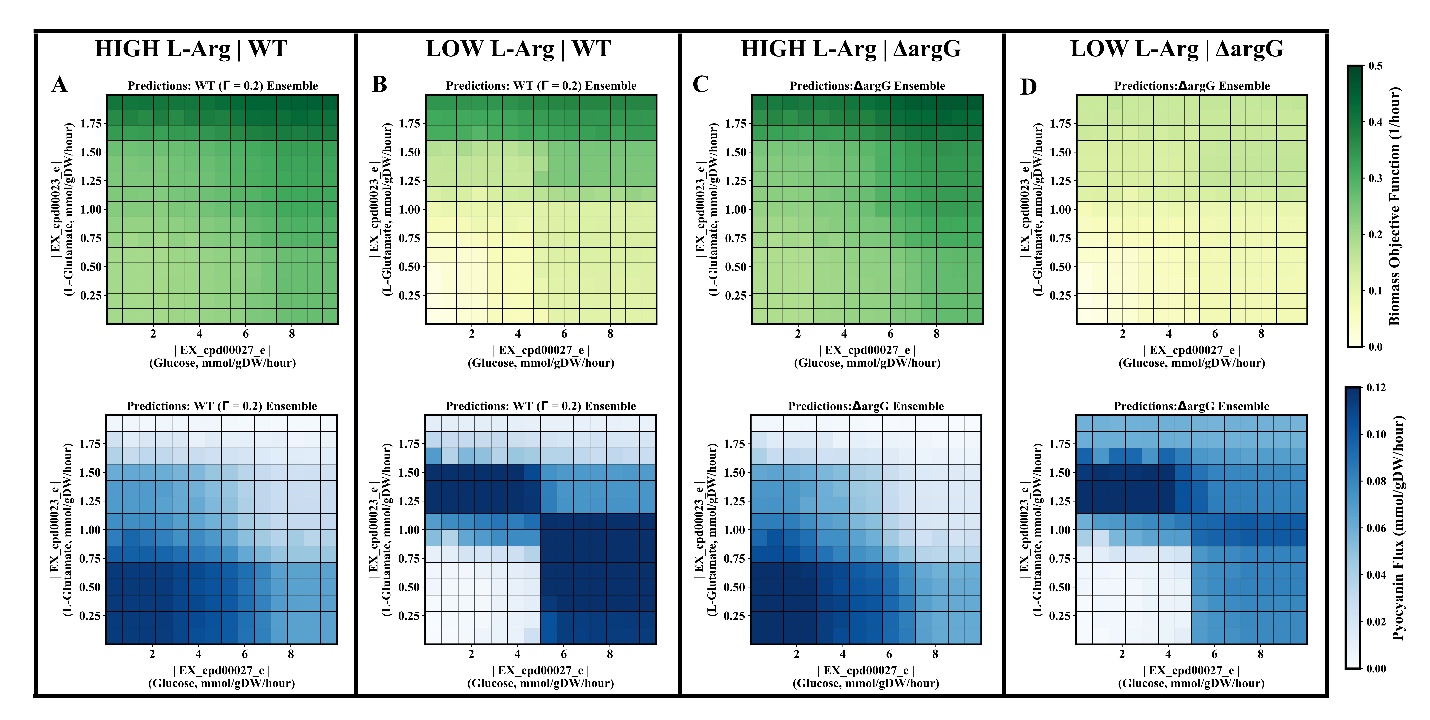

**Fig S.5**. **Ensemble coarse-graining of ΔargG metabolic behavior versus WT**. Heatmaps of biomass objective function and PCN flux predictions from (**A**, **B**) WT (Γ = 0.2) and (**C, D**) ΔargG ensembles at varying levels of L-arginine input flux. Glucose and L-glutamate inputs to each ensemble were varied with L-arginine input flux either set to -0.1 mmol/gDW/hour (LOW L-Arg) or -1.4 mmol/gDW/hour (HIGH L-Arg).

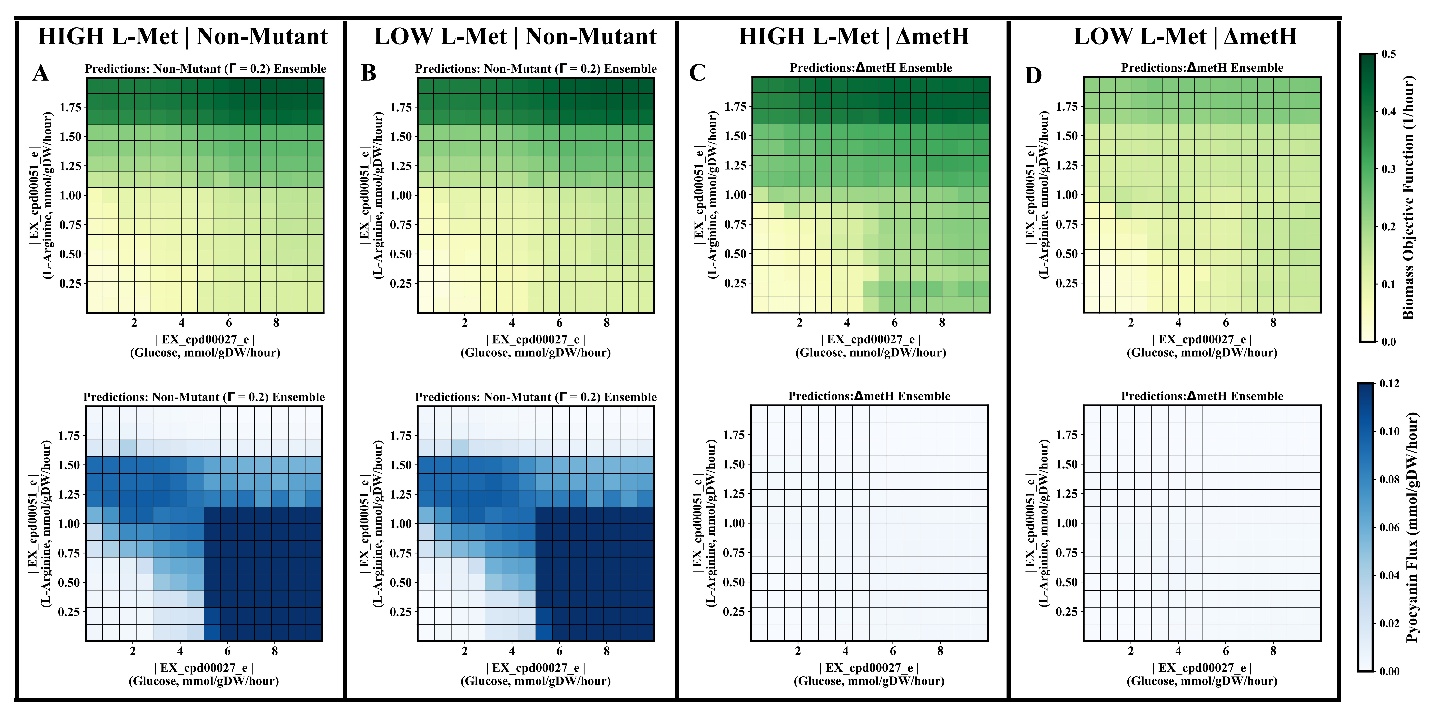

**Fig S.6**. **Ensemble coarse-graining of ΔmetH metabolic behavior versus non-mutant case.** Heatmaps of biomass objective function and PCN flux predictions from (**A, B**) non-mutant and (**C, D**) ΔmetH ensembles at varying levels of L-methionine input flux. The non-mutant ensemble was trained using an FBA dataset without any *in silico* gene deletion contexts and had glucose, L-arginine, and L-methionine as input features. Glucose and L-arginine inputs to ΔmetH and non-mutant ensembles were varied with L-methionine input flux either set to -0.04 mmol/gDW/hour (LOW L-Met) or -0.8 mmol/gDW/hour (HIGH L-Met).

### ***Appendix F****: Parameters for the genome-scale, metabolism-informed continuum models (GEM-CM).*

***F.1:*** *Spatial and temporal discretization.*

**Table S.8. GEM-CM continuum layer variables for simulation mesh discretization and timesteps.**

| **Variable** | **Value** | **Unit** |
| --- | --- | --- |
| **Plate Radius** | **1.74** | **cm** |
| **Space Discretization** | **0.002** | **cm** |
| **Simulation Duration** | **20** | **hour** |
| **Time Discretization** | **1** | **second** |
| **Ensemble Update Interval** | **5** | **minute** |

**Table S.8** summarizes the spatial mesh and time discretization used to construct and evaluate the GEM-CM continuum layer. The continuum environment depicts the swim plate assay as a one-dimensional radial coordinate grid covering the radius of a Corning 6-well dish. The spatial grid was constructed using mesh design procedures reported by previous investigations [3, 9]. Partial differential equations (PDE) for PA14 and metabolites were evaluated using the finite volume solver FiPy (v3.4.2.1) [10] over one second timesteps for a total of 20 simulated hours. At the start of each five minute time interval during the simulation, carbon source concentrations were passed to the ensemble model to obtain updated nutrient consumption, PCN production, and PA14 growth rates at each spatial mesh point. Nutrient concentrations in the continuum layer were reviewed each time after the finite volume solver returned the PDE solution. If any mesh spaces encountered depletion in any of the carbon sources before the next five minute interval, all rate values at those mesh location(s) were substituted with new ensemble model predictions.

***F.2****: Determining bacteria diffusion coefficient from macrocolony spreading velocity.*

The PDE expression for PA14 biomass in the GEM-CM continuum layer has notable resemblance to the Fisher-KPP equation [11, 12] in radial coordinates (**Eq. S3**) if reaction rate *‘g’* is substituted for Monod growth kinetics expression (**Eq. S4**).

$$\frac{\partial C_{B}}{\partial t}=\frac{D_{B}}{r}\frac{\partial}{\partial r}\left[ r\frac{\partial C_{B}}{\partial r} \right]+gC_{B}\left[ 1- C_{B} \right]$$

**(Eq. S3)**

$$g=\mu_{max}\frac{[C_{S}]}{k_{S}+[C_{S}]}$$

**(Eq. S4)**

The resulting propagation of the species *C_B_* has been described as a traveling wave moving with a stable speed that can be expressed using the diffusion coefficient and reaction rate. When examining **Eq. S3** at the leading edge of the macrocolony where biomass density (*C_B_*) is low, the order (*C_B_*)^2^ terms in **Eq. S3** become negligible and the remaining terms all match the PDE expression in **Eq. 4.** Thus, the Fisher-KPP equation’s traveling wave speed is also applicable to the GEM-CM PA14 population. Given that nutrient concentration (*C_S_*) near the macrocolony leading edge is also likely large relative to the half-velocity constant (*k_S_*), the reaction rate in the traveling wave velocity (**Eq. 2**) reduces to *µ_max_*. A representative value for *µ_max_* (0.76 hour^-1^) was estimated by passing the starting concentrations of glucose, L-arginine, and L-glutamate to the WT (Γ = 0.2) ensemble after values for the ensemble-continuum interface parameters were evaluated (discussed in following section). The set of bacteria motility coefficients used for 0.25% and 0.28% GEM-CM simulations are summarized in **Table S.9**; we assigned metabolites and PCN identical, constant diffusion coefficient values assuming that their transport was not affected by the water-swelled agar gel or molecular interactions.

**Table S.9. Diffusion coefficients for GEM-CM continuum layer.**

| **Variable** | **Value(s) (cm^2^/second)** | **Reference** |
| --- | --- | --- |
| PA14, 0.25% agar **^(a)^** | **1.49 E-6** | **This study** |
|  | **1.03 E-6** |  |
|  | **1.30 E-6** |  |
|  | **0.89 E-6** |  |
|  | **1.63 E-6** |  |
| PA14, 0.28% agar **^(a)^** | **0.50 E-6** | **This study** |
|  | **0.45 E-6** |  |
|  | **0.57 E-6** |  |
|  | **0.38 E-6** |  |
|  | **0.54 E-6** |  |
| Metabolites | **8.00 E-6** | **Cremer *et al.* 2019** |

**(a) Bacteria motility coefficients identified for each swim plate assay replicate using its measured macrocolony spreading velocity. A GEM-CM batch simulation was performed for each bacteria diffusivity value in a gel composition group with all other model parameters and initial conditions held constant.**

***F.3****: Determining parameters for metabolite concentration conversion to mass flux.*

In order to couple the ensemble and continuum layers, concentration fluctuations must be translated to ensemble input values that reflect an appropriate change to apparent mass flux. The relationship between extracellular concentration and metabolite flux defined by **Eq. 1** is shaped by the values of the maximum exchange flux (*v_max_*) and the half-velocity constant (*k_S_*). To identify suitable *v_max_* and *k_S_* values for each metabolite input in the ensemble, we scored the dynamic simulation predictions from candidates of parameter sets against liquid culture measurements collected by McGill *et al.* 2021 [13] for PA14 in a defined growth media with similar metabolite composition as M9-glucose. In place of the swim plate PDE system, ordinary differential equations (**Eq. S5-S7**) were defined to depict a well-mixed liquid culture.

$$\frac{dC_{B}\left( t \right)}{dt}=v_{biom}C_{B}\left( t \right)$$

**(Eq. S5)**

$$\frac{dC_{N,i}\left( t \right)}{dt}={-v}_{\max,i}\left( \frac{C_{N,i}}{k_{S,i}+C_{N,i}} \right)C_{B}\left( t \right)$$

**(Eq. S6)**

$$\frac{dC_{PCN}\left( t \right)}{dt}=v_{PCN}C_{B}\left( t \right)$$

**(Eq. S7)**

Initial metabolite concentrations and biomass density were matched to McGill *et al*. reports of starting liquid culture conditions. Dynamic simulations were also initiated with *v_max_* and *k_S_* parameter values (three *v_max_* and three *k_S_* in total) and the WT (Γ = 0.2) ensemble to predict reaction rate terms in **Eq. S5-S7**. Routines for updating reaction rates with new ensemble predictions were similar to routines described in *Appendix F.1*. After reaching the simulated time endpoint, predictions for each carbon source concentration (*C_N_*) at discrete timepoints were compared to experimental metabolite concentrations measured by McGill *et al*.; the normalized RMSE for each metabolite species was then calculated and added together for a final *loss function* output.

To minimize the loss function formed by the set of *v_max_* and *k_S_* parameter, we implemented Bayesian optimization procedures practiced by Khazaei *et al.* [14] to identify appropriate FBA constraints for dynamic metabolic models. The search space for amino acid *ν_max_* and *k_S_* were constrained to [-5, -0.5] mmol/gDW/hour and [1 – 50] µM, respectively. The search space for glucose *ν_max_* was extended to [-10, -0.5] mmol/gDW/hour while maintaining the same *k_S_* parameter constraints as was used for amino acids. First, the posterior distribution of the loss function was constructed from 10 randomly sampled candidate sets of *ν_max_* and *k_S_*. Bayesian optimization was then allowed to sequentially evaluate n = 50 new candidates to improve on the loss function minimum. The best candidate set of *ν_max_* and *k_S_* after 10 + 50 iterations was recorded and the entire optimization process starting at the posterior distribution construction was repeated a total of 50 times. A geometric mean for each parameter was calculated using the global top 10 candidates and set as the *ν_max_* and *k_S_* for all ensemble models (**Table S.10**). When *ν_max_* and *k_S_* had to be determined for carbon sources not originally included in the WT ensemble, Bayesian optimization was performed using a new ensemble trained on the foundation of glucose, L-glutamate, and L-arginine exchange reaction inputs as well as any additional carbon source(s) of interest.

**Table S.10. Monod kinetics parameters for coupling ensemble and continuum layers in GEM-CM.**

| **Metabolite** | ***v_max_***  **(mmol/gDW/hour)** | ***k_S_***  **(µmol/L)** |
| --- | --- | --- |
| **Glucose** | **9.86** | **17** |
| **L-Arginine** | **2.68** | 4 |
| **L-Glutamate** | **4.86** | **8** |
| **L-Methionine** | **0.83** | **11** |

### ***Appendix G****: Flux variability analysis for ΔmetH deletion.*

Flux variability analysis (FVA) [15] is a constraint-based analysis technique which can estimate the lower and upper mass flux bounds of iPau21 GEM reactions that would allow the objective function to reach a cutoff *fraction of the optimum*. The mass flux range between the lower and upper bounds of a given reaction can then be used to interpret whether the objective function is sensitive or insensitive to the reaction. We performed FVA on the GEM before and after *metH* (PA14_40670) deletion setting biomass flux as the primary objective and the fraction of the optimum to 90%. The complete FVA result summary can be accessed through this study’s GitHub repository, while **Table S.11** reports minimum and maximum flux bounds of select GEM reactions. Nutrient media exchange fluxes were maintained at their “high growth” condition settings (**Table S.6**) with the exception of L-methionine exchange which had its overall mass flux reduced in order to emphasize the effect that ΔmetH has on intracellular L-methionine supply.

**Table S.11. Flux variability analysis of select reactions in the iPau21 GEM***.*

| **Reaction** | **iPau21 Role** | **No Deletion (mmol/gDW/hour)**  **[minimum, maximum]** | **ΔmetH (mmol/gDW/hour)**  **[minimum, maximum]** |
| --- | --- | --- | --- |
| **PA14_Biomass** | **PA14 Growth** | **[0.81, 0.90]** | **[0.81, 0.90]** |
| **EX_cpd00027_e** | **Glucose exchange** | **[-10, 0]** | **[-10, 0]** |
| **EX_cpd00051_e** | **L-Arginine exchange** | **[-5, 0]** | **[-5, 0]** |
| **EX_cpd00023_e** | **L-Glutamate exchange** | **[-5, 0]** | **[-5, 0]** |
| **EX_cpd00060_e** | **L-Methionine exchange** | **[-0.10, 0]** | **[-0.10, -0.07]** |
| **rxn00693** | **Homocysteine** $\to$  **L-Methionine** | **[-0.03, 0.28]** | **[0, 0]** |
| **rxn00126** | **S-adenosylmethionine synthesis** | **[0.01, 0.21]** | **[0.01, 0.04]** |
| **rxn13896** | **Pyocyanin synthesis** | **[0, 0.20]** | **[0, 0.03]** |

Reflecting its gene-protein-reaction connection to *metH*, rxn00693 displayed total mass flux elimination in the knockout FVA. PA14 growth however, maintained similar minimum and maximum bounds across both deletion-absent and ΔmetH contexts suggesting that exchange flux constraints for “high growth” conditions could circumvent the growth-inhibiting effect of *metH* knockout. In contrast, L-methionine exchange flux range was notably reduced in ΔmetH compared to the deletion-absent case, which indicated that extracellular L-methionine has elevated importance for ΔmetH when trying to attain 90% fraction of optimum. This outcome and rxn00693 are connected by *metH* serving a critical role in the L-methionine biosynthesis KEGG pathway, and thus the ΔmetH context being more reliant on growth media L-methionine to supply intracellular demands. Disruption to rxn00693 also concurrently limited mass flux through S-adenosylmethionine (SAM) and PCN synthesis reactions. When paired with the mechanisms illustrated by **Fig 7**, the FVA results for these synthesis reactions support a L-methionine bottleneck in the *metH* knockout limiting SAM production and major reactions that involve SAM, such as PCN synthesis [16].

### ***Appendix H****: GEM-CM captures PA14 growth and PCN production fluctuations for slower spreading macrocolonies.*

Changes to an external environment’s physical properties have been shown to impact the displacement and navigation of motile *P. aeruginosa* [3, 17]. To evaluate how GEM-CM predicts macrocolony development and PCN production in motility-altered PA14 populations, we prepared and observed swim plate assays possessing gels with higher (0.28% w/v) agar density. Based on results reported for *Escherichia coli* in variable agar weight percentages, we expected PA14 macrocolonies to have slower spreading in the 0.28% swim agar due to smaller pore sizes in the agar network restricting the cell’s swimming pattern, and consequently, lowering overall mean displacement [18]. Timelapse imaging confirmed PA14 macrocolonies in 0.28% w/v agar had ~2-fold decrease to spreading velocity resulting in reduced macrocolony size at t = 20 hours (**Fig S.7A-B**). Macrocolony spreading velocities and Fisher-KPP wave velocity equation (**Eq. 2**) were used to determine new *D_B_* values (0.3 – 0.6 E-06 cm^2^/s) for depicting slower PA14 motility in the GEM-CM model. Simulations of PA14 in 0.28% agar still retained the ensemble model along with *v_max_* and *k_S_* parameters used in the 0.25% condition, thus assuming that growth was not affected by the increased agar density.

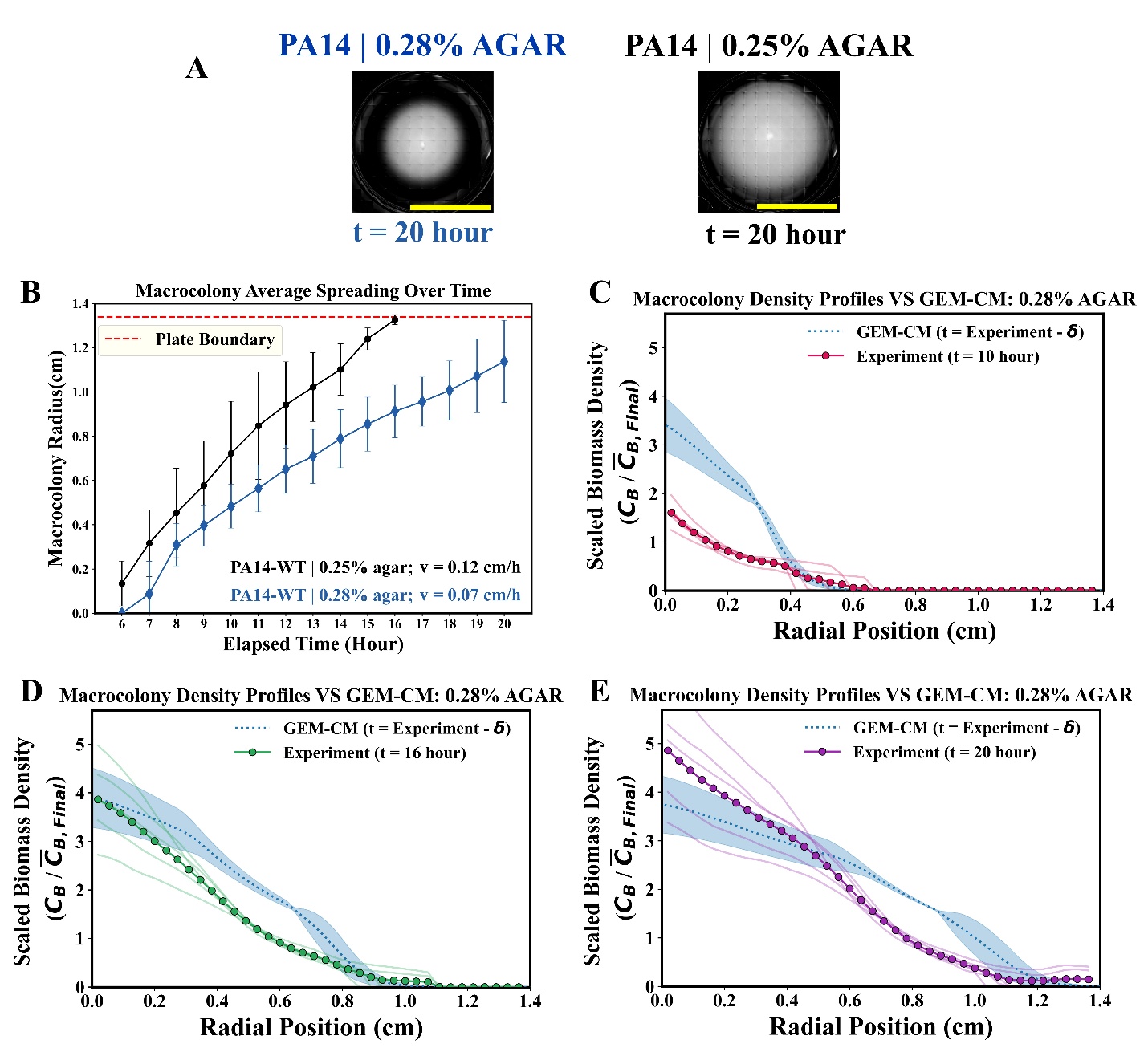

**Fig S.7**. **PA14 swim plate assays and simulations in 0.28% agar.** (**A**) Tiled brightfield images of PA14 macrocolonies (opaque white regions) in 0.28% versus 0.25% swim agar at end of timelapse observation. Scale bar = 2 cm. (**B**) Average macrocolony radii versus time and mean spreading velocity estimations of 0.25% (transferred from **Figure 3**, black dots) and 0.28% (n = 5, blue diamond) agar swim plates; error bars = 1 standard deviation. Red dashed line denoting the distance to the impermeable well dish wall. (**C - E**) Experimental and GEM-CM predicted biomass density profiles at discrete time points scaled to each group’s average endpoint biomass density. GEM-CM density profile predictions are adjusted by 𝛅 (two hours). Group average and sample replicate density profile visualization follow the schemes from **Fig 3.**

After adjusting simulated time by the estimated lag-phase from 0.25% agar swim plate measurements, GEM-CM predictions for biomass density profiles in 0.28% agar largely returned the trends of over-prediction at early time points, fair agreement with experimental macrocolony spreading distance, and improving accuracy over longer timescales (**Figure S.7C-E**). By the assay endpoint, the average GEM-CM predictions had an R^2^ = 0.87 against measured 0.28% agar swim plate profiles at t = 20 hour. These comparable model prediction performances between 0.25% and 0.28% agar conditions further supported the applicability of spreading velocity and Fisher-KPP wave speed to estimate appropriate *D_B_* parameter values. Still, the presence of high relative PA14 biomass density near the center of 0.28% agar macrocolonies motivated us to examine GEM-CM spatial growth and PCN production rate dynamics in order to extract additional model framework context around the effects of constrained bacterial motility on population-level outcomes.

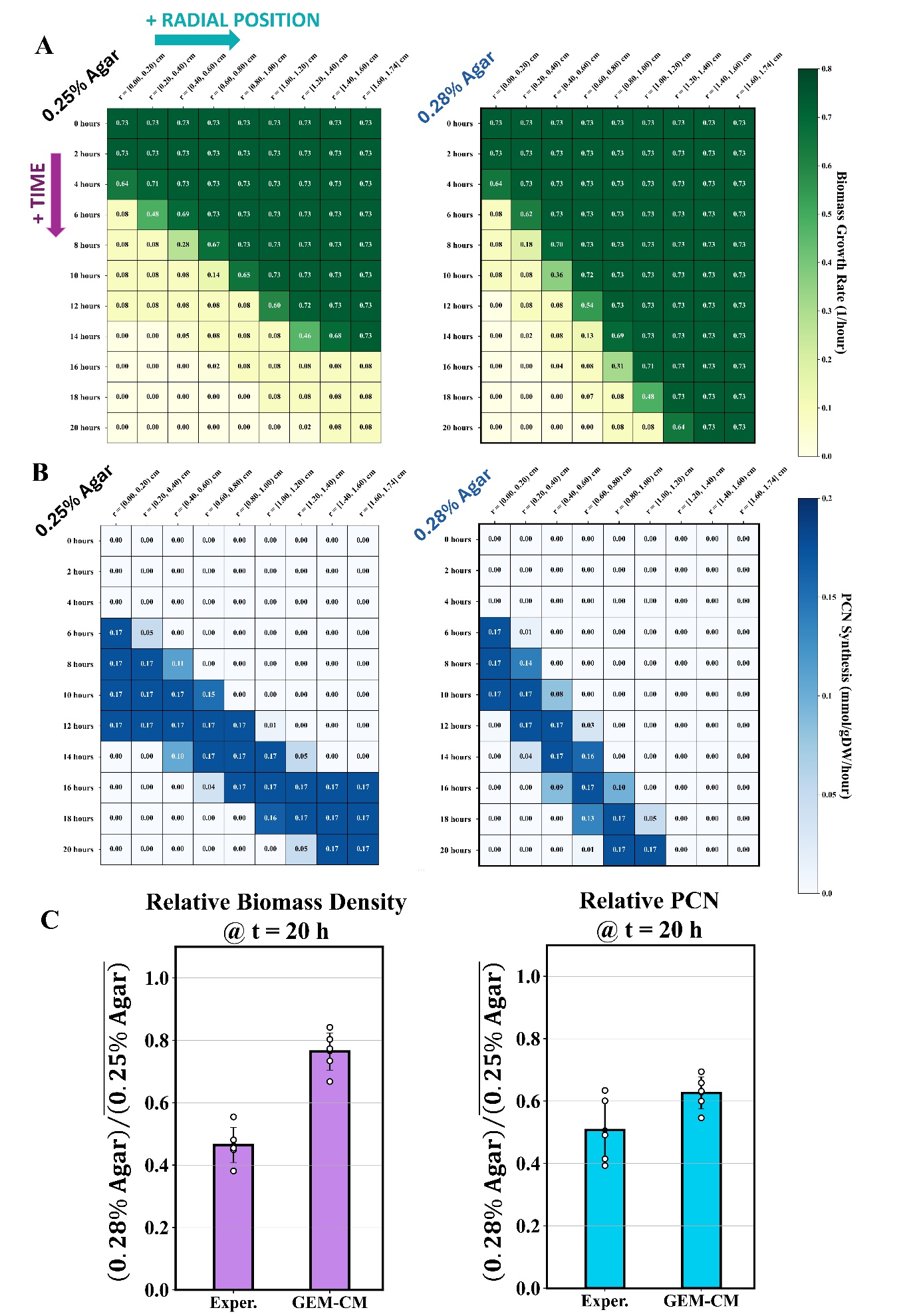

**Fig S.8.** **Effects of reduced PA14 motility on biomass growth and PCN production in GEM-CM predictions**. (**A**) 2-D heatmaps of predicted growth rate over space and simulated time for a 0.25% agar swim plate simulation (*D_B_* = 1.5 E-06 cm^2^/s) and a 0.28% agar swim plate simulation (*D_B_* = 0.5 E-06 cm^2^/s). Cell values denote the average growth rate within a specified radial position range and time point. Radial position increases left-to-right along the heat map and simulated time increases going from top-to-bottom. (**B**) 2-D heat maps for PCN production rates from the same GEM-CM simulation run as their growth rate counterpart in **A**. (**C**) Biomass density and PCN concentrations from 0.28% agar experiments and simulations relative to their mean quantities from the 0.25% agar condition. Scatter plot markers overlaying bar graphs represent each experiment or GEM-CM replicate.

The heatmaps of predicted growth rates over the GEM-CM simulated space (**Fig S.8A**) additionally convey the overall quantity of nutrients remaining in a localized swim plate area, as illustrated by the top row of each heat map (*t_sim_* = 0 hour) monotonically having the maximum growth rate feasible given the starting carbon source concentrations. In the 0.25% agar condition, PA14 was able to spread the full radial distance of swim plate model from the r = 0 inoculation point, and the macrocolony consumed nearly all available carbon sources over the course of the simulation. In contrast, decreased PA14 motility in the 0.28% agar condition impeded PA14 from reaching the outer edge of the swim plate model and left the carbon sources in that region (right-most column) inaccessible. Motility-constrained macrocolonies also had fewer total swim plate regions with active PCN production during their simulation (**Fig S.8B**), further underlining that slower macrocolony spreading limits the development and metabolic activity a population can accomplish within a discrete time span. Together, biomass growth and PCN production spatiotemporal trends in GEM-CM simulations yielded lower average biomass density and PCN for 0.28% agar conditions relative to 0.25% agar (**Fig S.8C**), which also agreed with assay measurements. However, biomass density reduction between 0.28% and 0.25% agar was more drastic in experiments than in simulation predictions, which may suggest that *in vitro* PA14 growth is affected by other local factors separate from nutrient availability. Future investigations could consider alternative cell growth descriptions, like the logistic growth equation, to more fully examine growth behavior in these macrocolonies.
